# A neural circuit mechanism for abstract free choice

**DOI:** 10.64898/2026.05.29.728548

**Authors:** Jake Gavenas, Tian Lan, Aaron Schurger, Uri Maoz

**Affiliations:** Department of Neurosurgery, Cedars-Sinai Medical Center, Los Angeles, CA, USA; Institute for Interdisciplinary Brain and Behavioral Sciences, Chapman University, Orange, CA, USA; Schmid College of Science and Technology, Chapman University, Orange, CA, USA; Crean College of Health and Behavioral Sciences, Chapman University, Orange, CA, USA; Fowler School of Engineering, Chapman University, Orange, CA, USA; INSERM U992, Cognitive Neuroimaging Unit, NeuroSpin Center, Gif sur Yvette 91191, France; Commissariat à l’Energie Atomique, Direction des Sciences du Vivant, I2BM, NeuroSpin Center, Gif sur Yvette 91191, France; Division of Biology and Biological Engineering, California Institute of Technology, Pasadena, CA, USA; Anderson School of Management, University of California, Los Angeles, CA, USA

**Keywords:** Free choice, abstract decision-making, goal switching, attractor dynamics, EEG, winner-take-all circuit, hierarchical processing

## Abstract

Making abstract free choices, such as arbitrary decisions between comparable options formed independently of specific motor plans, is central to everyday behavior. Yet, the neural mechanisms underlying free choice remain poorly understood. Using a delayed-response paradigm (N=80), we dissociated abstract goal selection (color choice) from later motor response mapping. We found that freely chosen goals elicit slower responses and lower goal-switching costs than instructed goals. We developed a hierarchical attractor network model in which symmetry-breaking attractor dynamics drive free goal selection, and conjunctive representations translate these abstract goals into motor output. Freely chosen goals converged on less extreme fixed points than instructed ones, accounting for their slower, more variable responses. Consistent with our model, human EEG activity (N=30) sequentially encoded goal, conjunctive, and response representations, with conjunctive representations specifically coupled to behavioral variability. Our findings offer plausible circuit-level mechanisms for how abstract free choices are made and translated into action.

**Highlights:**

1. Free-instructed differences persist when goal selection and response mapping are dissociated
2. Abstract free choices elicit reduced goal-switching costs relative to instructed ones
3. Symmetry-breaking attractor dynamics explain behavioral signatures of free choice
4. EEG signals sequentially encode abstract goal, conjunctive, and response representations
5. Conjunctive representations, but not goal or response, were coupled to behavior

## Introduction

The ability to make arbitrary decisions between functionally equivalent options is a fundamental behavioral capacity. Consider approaching an obstacle while driving on an open road: you could swerve left or right but decisively committing to one option and then acting on your choice is paramount. Such choices are ubiquitous in everyday life: we frequently choose between comparable alternatives, such as where to go for dinner or what brand of toothpaste to buy, before acting on our decision. Despite their ubiquity, however, the neurocognitive mechanisms underlying “free choices”—decisions not immediately driven by instruction, memory, or value—remain poorly understood.

A canonical behavioral trait of free choice is that response times for freely chosen actions are slower than for instructed ones (aka fixed or forced choice)^1–6^. It has been suggested that the need to engage additional decision circuits in the case of free choice, but not when instructed, drives such reaction time differences ^1,5^. Studies finding that free choices activate frontoparietal^7–15^ and striatal^8,9^ regions have been considered evidence in favor of this hypothesis. However, more recent behavioral and neuroimaging studies support common underlying neural representations for free and instructed choice^2,3,16^; for example, similar fMRI representations were found for choice *content* across free and instructed contexts^17,17,18^. If free and instructed choices rely on the same underlying representations, what drives the distinct behavioral signatures of free choice? We suggest the answer lies not in separate cognitive steps, but in the intrinsic dynamics of the neural networks computing these choices.

Puzzlingly, the *content* of free choices can be predicted with significantly above-chance accuracy several seconds before the decisions are expressed^7,12,13,19^, or even before decision alternatives have been presented^9,20^. It is possible that such seemingly paradoxical findings are due to attractor network dynamics. Specifically, Rolls & Deco^21^ were sometimes able to predict choice content using pre-stimulus activity in an attractor network model of perceptual decision-making that lacked contextual factors like choice history. Instead, spontaneous fluctuations in pre-stimulus activity biased the network when decision nodes received symmetric input — as may be the case in free choice — allowing the initial state to partially determine the outcome before evidence arrives. Some have suggested such a mechanism could explain free choice behavior^9,20^. But this hypothesis has not been empirically tested, in part because standard free-choice paradigms that operate at the level of immediate action selection conflate goal selection with motor preparation (i.e., the cue to make a choice is also the cue to execute the action), making it difficult to isolate the dynamics of the decision itself.

In the present study, we investigated the neurocognitive mechanisms underlying abstract free choice and their translation into motor output using a variable-delay paradigm. Crucially, our paradigm enabled us to test whether symmetry-breaking attractor dynamics could explain behavioral signatures of free choice. First, participants were either instructed or freely chose an abstract color (blue or green). After a variable delay, they were presented with a response map matching the two colors to a left/right button press. By separating goal selection from response mapping, we directly tested whether the observed response-time differences between free and instructed conditions were due to an additional cognitive step in the decision-making process for free choice, or whether free choices were encoded differently from instructed actions within the same neural networks. We additionally occasionally cued participants to ignore prior goals and respond in the left or right direction upon receiving the response mapping, to characterize decision commitment beyond just reaction time^22^.

To map our results onto specific circuit-level mechanisms, we developed a hierarchical attractor network model^22,23^ that accounted for our behavioral results through a combination of three key mechanisms: (1) symmetry-breaking winner-take-all dynamics for free choices^21,24^; (2) temporary suppression of motor output through a direct inhibitory pathway^25,26^; and (3) conjunctive representations that integrate abstract choice and motor mapping to serve as an intermediary before motor output^27–29^—a component we directly tested using EEG in a second study that replicated our behavioral findings. Altogether, our results support symmetry-breaking attractor dynamics and conjunctive representations that integrate abstract goals and sensory information as circuit-level mechanisms for how abstract free choices are made and then translated to motor output.

## Results

Participants (N=80; see Methods) completed a variable stimulus-onset-asynchrony (SOA) abstract choice task (Fig 1). The initial cue comprised two discs (each either blue or green) above and below a fixation cross. If both were the same color (67% of the trials), participants had to press in the direction of that color when presented with the motor mapping (Instructed Context). If the discs were different colors (33% of the trials), participants had to immediately select either blue or green and then press in that direction when presented with the motor mapping (Free Context; Fig. 1A). We refer to the color instructed or chosen as the Goal and whether the goal was freely chosen or instructed as the Context.

**Figure 1.**
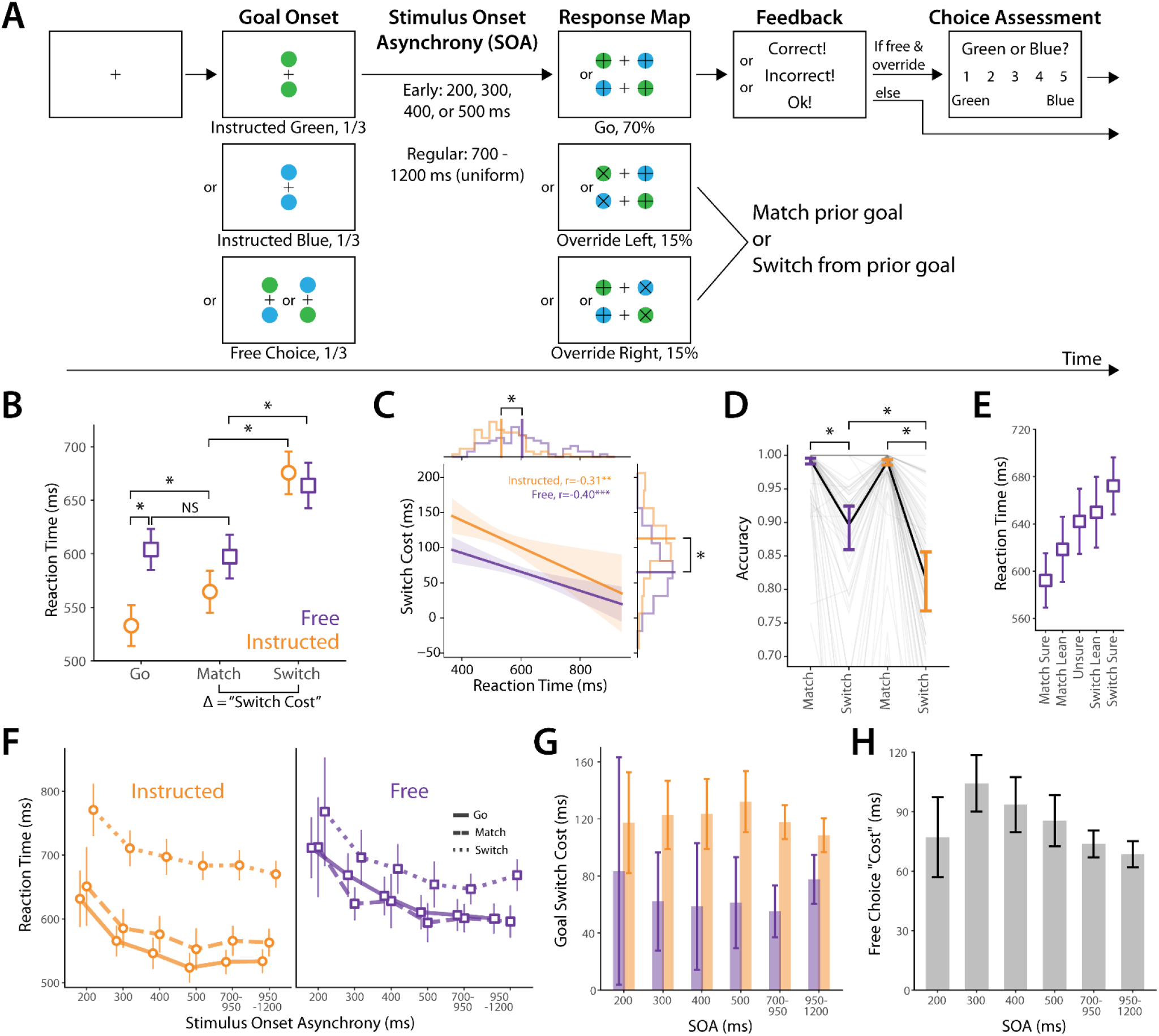
Task and behavioral results. **(A)** Abstract free choice task. From left to right: Participants instructed to select a goal of blue (top), green (middle) or to freely pick blue or green (bottom). Then, after a variable stimulus onset asynchrony, they were presented with the motor mapping of blue/green goal to left/right response or vice versa (top, Go trials). On Override trials, an X instructed participants to forego their prior goal and press in the direction of the X (middle and bottom). Participants were then given feedback (“OK!” on Free trials, where decision correctness was not defined, and “Correct!” or “Incorrect” on all other trials based on their performance). On Free Override trials, participants then reported their confidence in their decision (see Methods). This enabled us to know whether their button press entailed a Match or a Switch from their initial choice. **(B)** Main RT results for regular SOA range (700-1200 ms; mean & 95% confidence intervals, obtained from LME marginal estimates). (**C)** Correlation of RTs from Go trials with goal switch costs estimated from override trials (regular SOA range) alongside histograms of individual subjects’ values (estimated as random intercepts/effects from LME model; thick solid lines perpendicular to the axes reflect mean values). (**D)** Override trial accuracy (means & 95% confidence intervals; logistic mixed effects; gray lines = individual participants). **(E)** Reaction times on free choice override trials as a function of subjective goal commitment (means & 95% confidence intervals). **(F)** RTs in all conditions as a function of SOA (means & 95% confidence intervals). **(G)** Switch costs as a function of SOA (means & 95% confidence intervals). **(H)** Free choice cost (estimated difference between Free and Instructed Go trial RTs; means & 95% confidence intervals) as a function of SOA.

After a variable SOA (700-1200 ms uniform for “regular” SOAs (70% of trials), or 200, 300, 400, or 500 ms within the “early” SOA (each equally likely, together 30% of trials); stimuli remained onscreen during the delay), participants were presented with the response map, consisting of a blue and a green disc on either side of the fixation cross (Map; blue-green or green-blue; Fig. 1A). 70% were Go trials, with a + sign on both discs, which indicated that participants should press in the direction mapped to their instructed or chosen color. We also wanted to quantify decision commitment in addition to reaction time (RT) alone. Thus, on 30% of trials, one of the discs had an X over it (Fig. 1A), indicating that the participant should press in that direction *regardless of their color selection* (Override trials). The response demanded by override cues could either be consistent (Match Trials) or inconsistent (Switch Trials) with their prior goal (Match and Switch trials respectively, with Go, Match, or Switch collectively referred to as Map Type). Both trial types involved incorporating the override cue, but only Switch trials involved delivering a different response than they otherwise would have. Thus, comparing these trials allowed us to quantify decision commitment.

After responding, participants were given feedback about whether their response was correct or not, on Instructed or Free Override trials, or the feedback “Ok”, on Free Go trials. On Free Override trials only, we next asked participants what color they had selected before receiving the override cue using a 5-point Likert scale (Decided Blue, Tended Blue, Undecided, Tended Green, Decided Green). Across participants, Undecided and Tending responses were infrequent relative to Decided (mean percentages±SEM, Decided=66.7±0.04%, Tending=32.1±0.03%, Undecided=23.2±0.03%), so we omitted Undecided trials and grouped Tending with Decided trials of the corresponding type in subsequent analyses unless otherwise indicated. After removing reaction time (RT) outliers and incorrect trials (see Methods) we were left with a total of 371±32 trials per participant out of a total scheduled 400 (range: 287-397).

### Free choice produced intact but diminished signatures of goal commitment

We first tested whether previously reported response slowing during free-choice trials emerged because participants had to also make decisions on those trials. We found that even when goal selection and response mapping were separated by a SOA of 700 ms or more, Free Go trials had significantly longer RTs than Instructed Go trials (mean [95% CI]; 604.1 [585.3,623.0] ms vs 533.0 [514.3, 551.7] ms; T(19439.3)=29.8, p<0.001; LME post-hoc tests with Tukey correction; Fig. 1B). SOA did not significantly affect Go trial RTs within this window (F(1,14031.3)=0.094, p=0.76, LME model with SOA and Context as factors), nor did it interact with Context (F(1,14031.4)=2.40, p=0.12, same LME as above), indicating that participants completed their decision making by the time the Response Map was presented. To make sure that the results are not due to a fraction of trials with especially long RTs we repeated this analysis for only the fastest 50% of RTs in each condition. The pattern of the results stayed the same (Fig S1A). Similarly, the results were not due to including trials wherein participants were only leaning towards rather than fully decided on their goal (Fig. S1B). The RT difference we observe (71.1 ms) is commensurate with recent studies that compare free and instructed choices^3,5^. This result thus provides evidence against the hypothesis that longer RTs for free choices reflect additional decision-related processing. Rather, it supports the hypothesis that freely chosen goals are encoded differently from instructed ones.

What is the nature of this differential encoding? We next investigated whether free decisions elicited qualitatively different behavioral effects from instructions, or similar effects with diminished magnitude. We first considered switching costs, defined as the cost specific to a response that switched from the action consistent with the original color choice to one inconsistent with it. We operationalized switching costs as the RT difference between Switch and Match trials, because these trials both contained an override cue, but only Switch trials entailed responding inconsistently with participants’ prior goal. Switching costs were significant in both Free (Free-Match: 597.4 [577.2, 617.6] ms vs Free-Switch: 663.8 [642.9, 684.7] ms; T(19441.6)=10.2, p<0.001; LME post-hoc test with Tukey correction) and Instructed contexts (Instructed-Match: 564.7 [545.3, 584.0] ms vs Instructed-Switch: 675.7 [656.1, 695.4] ms; T(19440.3)=24.6, p<0.001; same LME as above; Fig 1B). Switching costs were reliably larger for Instructed compared to Free conditions (113.2 ms vs 65.2 ms, W=455, p<0.001, Wilcoxon sign-rank test; Fig 1C) consistent with Free choice recruiting similar decision mechanisms as instructed choice, though to a lesser degree. Additionally, and critically, across participants switch costs negatively correlated with reaction times within the same context (Instructed r=-0.31, p=0.005; Free r=-0.40, p<0.001; linear regression on subject-specific mean RT and switch costs; Fig 1C) meaning faster responders exhibited higher switch costs. This indicates that, rather than two isolated effects, the longer reaction times for free choices and the smaller switching costs for these choices may be driven by the same latent cognitive variable—the strength of the commitment to the initial color goal.

We next investigated whether accuracy on override trials depended on the cue’s consistency with participants’ prior goal. We found that accuracy was lower for Switch than Match in both contexts (Free Match: 99.3%, Free Switch: 89.6%, Instructed Match: 99.0%, Instructed Switch: 81.6%; logistic mixed effects; Free Match vs Free Switch Z=9.9,p<0.001; Instructed Match vs Instructed Switch Z=17.7, p<0.001; logistic mixed effects post-hoc tests with Tukey correction; Fig. 1D). This suggests that, in addition to slower RTs on Switch trials, commitment to an initial goal was sometimes strong enough to prevent switching entirely. Futhermore, Instructed Switch trials had significantly lower accuracy than Free Switch trials (Z=5.3, p<0.001; same logistic mixed effects model as above). This provides converging evidence, alongside our reaction time data, that goal commitment remains intact but is diminished when comparing Free and Instructed contexts.

To assess whether participants had introspective access to their decision states, we mapped participants’ reported decision states (decided blue, lean blue, undecided, lean green, decided green; Fig 1A) onto the actual override trial type to construct a graded measure of decision certainty (sure match, lean match, unsure, lean switch, sure switch). We found that RTs scaled with their decision confidence (F(4,2008.9)=30.4, p<0.001; LME; Fig 1E). In our task, these ratings were provided after the motor response, hence this scaling could reflect either direct metacognitive access to the pre-cue decision state, or a post-hoc inference based on the cognitive effort required to execute the override response. Regardless of the specific metacognitive mechanism, the continuous relationship between RTs and subjective reports provides further evidence for a variable, graded state of goal commitment during free choice.

### Goal commitment emerges rapidly but more slowly for free than instructed choices

We next investigated whether we could track the emergence of goal commitment by investigating how the overall RT and accuracy effects we observed above break down as a function of SOA. We observed a decrease-then-plateau pattern of RTs for all conditions (Go, Match, Switch; Fig 1F). Unlike the regular SOA period, we found that RT significantly decreased as SOA increased for SOAs of 500 ms and below (F(1,8210.6)=121.1,p<0.001, LME analysis). SOA did not interact with other factors in this period (SOA*Context: F(1,8203.7)=0.35,p=0.55; SOA*Map: F(2,8203.3)=0.98, p=0.37; same LME as above), indicating that decreasing RTs were similar for all conditions.

This decrease-then-plateau RT pattern for Go and Match conditions makes sense based on increasing goal commitment over time. But one might expect the opposite behavior in Switch trials—an increase in RT as goal commitment increased—because a more deeply entrenched goal should take longer to override. However, because participants do not yet know the required motor mapping during the delay, this overall decrease-then-plateau pattern likely reflects an increase in general temporal readiness or non-specific action arousal (e.g., anticipating the imminent arrival of the map), rather than the preparation of a specific motor plan^30^. This general readiness effect may mask other goal-commitment effects. Thus, to investigate goal commitment separately from motor preparation, we investigated switch costs separately for each early SOA and in two bins within the regular SOA period (700-950 ms and 950-1200 ms). We found that switch costs were present and significant for all SOAs with similar magnitudes (Fig 1G; p < 0.001, except Free context 200, 300, and 400 ms SOAs, with p = 0.045, 0.001, and 0.010, respectively; LME analysis), indicating that goal commitment emerges rapidly following goal onset.

To investigate how the dynamics of goal commitment differed between Free and Instructed contexts, we investigated how RT differences between Free and Instructed Go trials varied with SOA (Fig 1H). RT differences were significant for all SOAs (all p < 0.001), with differences peaking at SOAs of 300 before decreasing to a stable plateau (see statistics below). The interaction between Context and SOA was significant when comparing the 200 and 300 ms bins (T(2355.9)=2.04, p=0.042, LME), indicating that the Free-Instructed RT gap was still growing within this period. That interaction term was not significant when comparing the 300 and 400 ms SOAs (T(3343.2)=-0.61, p=0.55), trending when comparing the 300 and 500 ms SOAs (T(3422.0)=-1.75, p=0.08), and significant when comparing the 300 and regular (700-1200 ms) SOAs (T(15734.2)=-4.24, p<0.001). Together with our earlier finding that SOA does not interact with Context during the regular SOA period, these results suggest that freely chosen goals require additional time to reach full commitment relative to instructed goals, but that they generally reach full commitment by the regular SOA period (Fig 1H). Participants’ accuracy on Switch trials (ratio of trials in which they were able to comply with the override cue when it conflicted with the motor mapping for their original color goal) showed a similar pattern. Instructed Switch accuracy fell below 100% for all SOAs, whereas Free Switch accuracy remained near ceiling until 400 ms SOA (Fig S2), converging with the RT evidence that freely chosen goals reach full commitment more slowly than instructed ones. Overall, these results suggest the dynamics of goal commitment are similar between Free and Instructed goals, but that full commitment to Freely chosen goals develops more slowly than Instructed ones.

### A Hierarchical Attractor Network Reproduces Free Choice Behavioral Signatures

To investigate the circuit-level basis of goal commitment differences between Free and Instructed contexts, we developed a hierarchical attractor network model ^22,23^ that integrated an abstract color goal with a later-presented response map to generate an appropriate action (Fig. 2A; see Methods for details). The network consists of a series of winner-take-all circuits ^24,31^ encoding the abstract goal; the response map; the override direction, if present; a conjunctive sensorimotor representation that integrates goal and map information ^27,29^; and a motor response. When devising this network, we strove for the simplest one that could well reproduce key elements of our empirical data.

**Figure 2.**
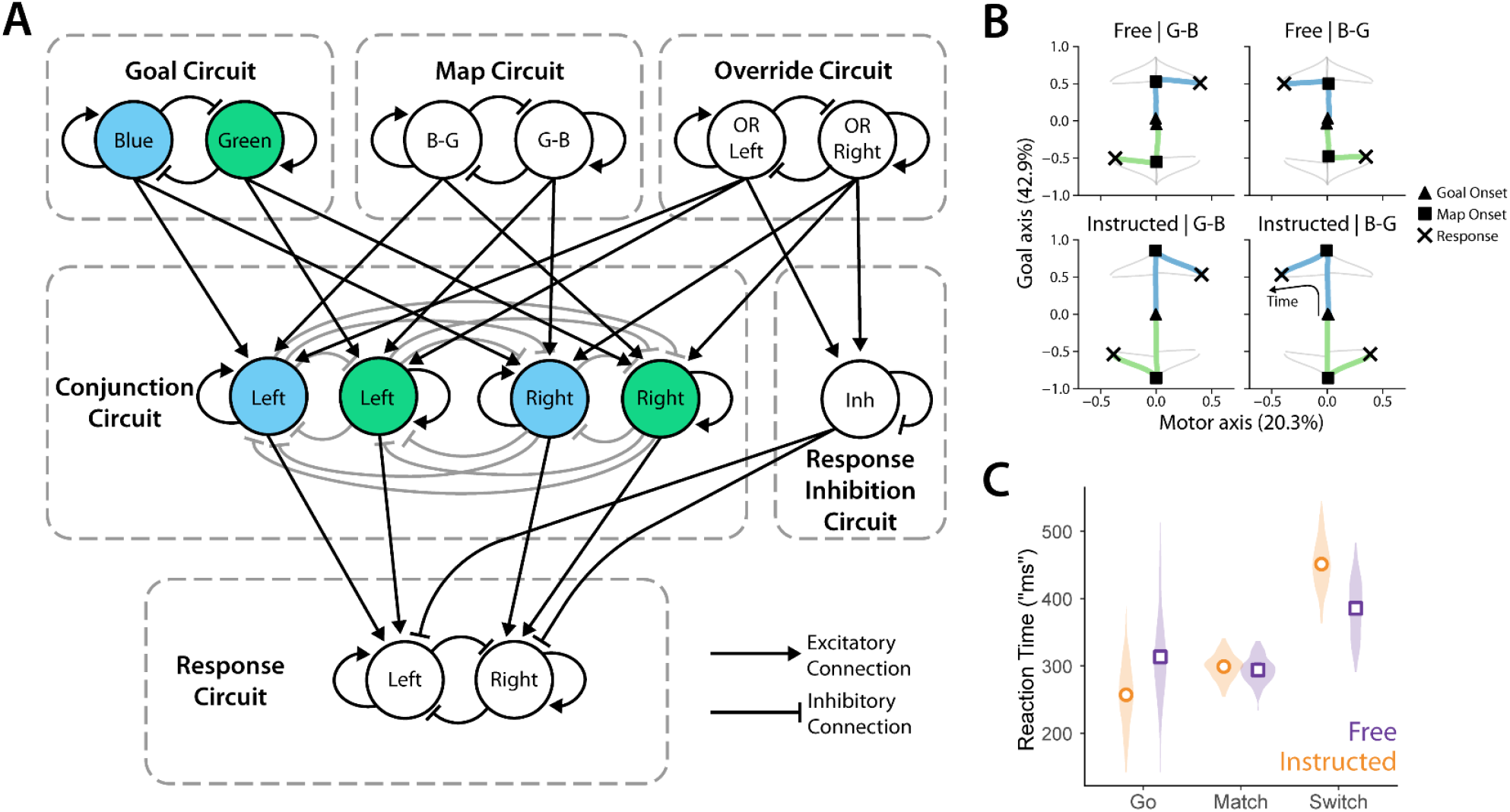
Hierarchical attractor network model for abstract free choice and goal switching. **(A)** Network diagram. The network comprises five winner-take-all (WTA) circuits and one direct inhibitory pathway, corresponding to (1) the blue/green goal circuit; (2) the map circuit, encoding left/right as blue/green or green/blue; (3) the override circuit, which encodes which side (if any) contains an override cue; (4) a conjunction circuit, integrating the abstract goal, the response map, and the potential override cue to enable action selection, and (5) the response circuit, which directly triggers left or right response once activity reaches a threshold. Additionally, if activated, the override circuit temporarily suppresses both motor responses via an inhibition circuit. Connections ending with an arrow (flat cap) are excitatory (inhibitory). **(B)** Projections of trial-averaged simulated activity onto orthogonal Goal and Motor axes. Activities shown are state-space trajectories averaged across individual trial runs and grouped by context (Free or Instructed) and map (B-G or G-B). Corresponding trajectories are in blue/green. Grey lines indicate other trajectories to visualize the overall structure of the state-space dynamics. Percentages in parentheses refers to the overall variability associated with these manually defined directions. **(C)** Violin plots of simulated RTs with means (estimated with LME) showing the distribution of all simulated trials.

The circuit operates as follows. On Instructed trials, one goal node receives preferentially stronger input. On Free trials, both nodes receive symmetric input, and noise drives the network to commit to a goal through winner-take-all dynamics (see Methods). In both cases, the settled goal representation primes the Conjunction Circuit to respond to the incoming Response Map in a goal-consistent manner (Fig. 2B; Fig. S3). To clarify these dynamics, we projected activity from Go trials onto manually defined orthogonal Goal and Response axes (see Methods). The model shows clear preparatory dynamics akin to those observed in delayed-response tasks ^30,32^. In such tasks, activity first builds along a preparatory axis during the delay before transitioning to an orthogonal response axis at movement onset (at Response Map onset in our case). On Go trials these dynamics proceed without interruption (Fig. S3A, B), but on Override trials activation of the response inhibition circuit leads to slower RTs (Fig S3C, D). Within override trials, Match trials exhibit relatively faster RTs because the direction of the override cue is consistent with activity in the conjunction circuit, leading to similar dynamics to Go trials (Fig S3A-C). However, Switch trials exhibit relatively slower RTs because the direction of the override cue is inconsistent with activity in the conjunction circuit, leading to disrupted conjunction circuit activity compared to Go trials (Fig S3D). Together, these dynamics reproduce our key empirical behavioral patterns in simulated RTs: Free Go trials were slower than Instructed Go trials, and Switch costs were larger for Instructed than Free contexts (Fig. 2C; compare Fig. 1B, G).

### Symmetry-Breaking Attractor Dynamics Predict Behavioral Signatures of Free Choice

What specific aspects of our model allow it to reproduce the behavioral signatures of free choice? To investigate this question, we considered the attractor dynamics within the circuit where Free and Instructed goal selection occurs (Fig 2A top left). In this circuit, each node self-excites and inhibits the other, leading to classic winner-take-all (WTA) dynamics^24,31^. We conducted vector field and fixed-point analyses of the goal circuit for the instructed and free conditions (Fig 3A, B), where black arrows reflect state-dependent gradients and colored trajectories reflect simulated dynamics. Red Xs mark stable fixed points, where trajectories converge and activity no longer evolves.

**Figure 3.**
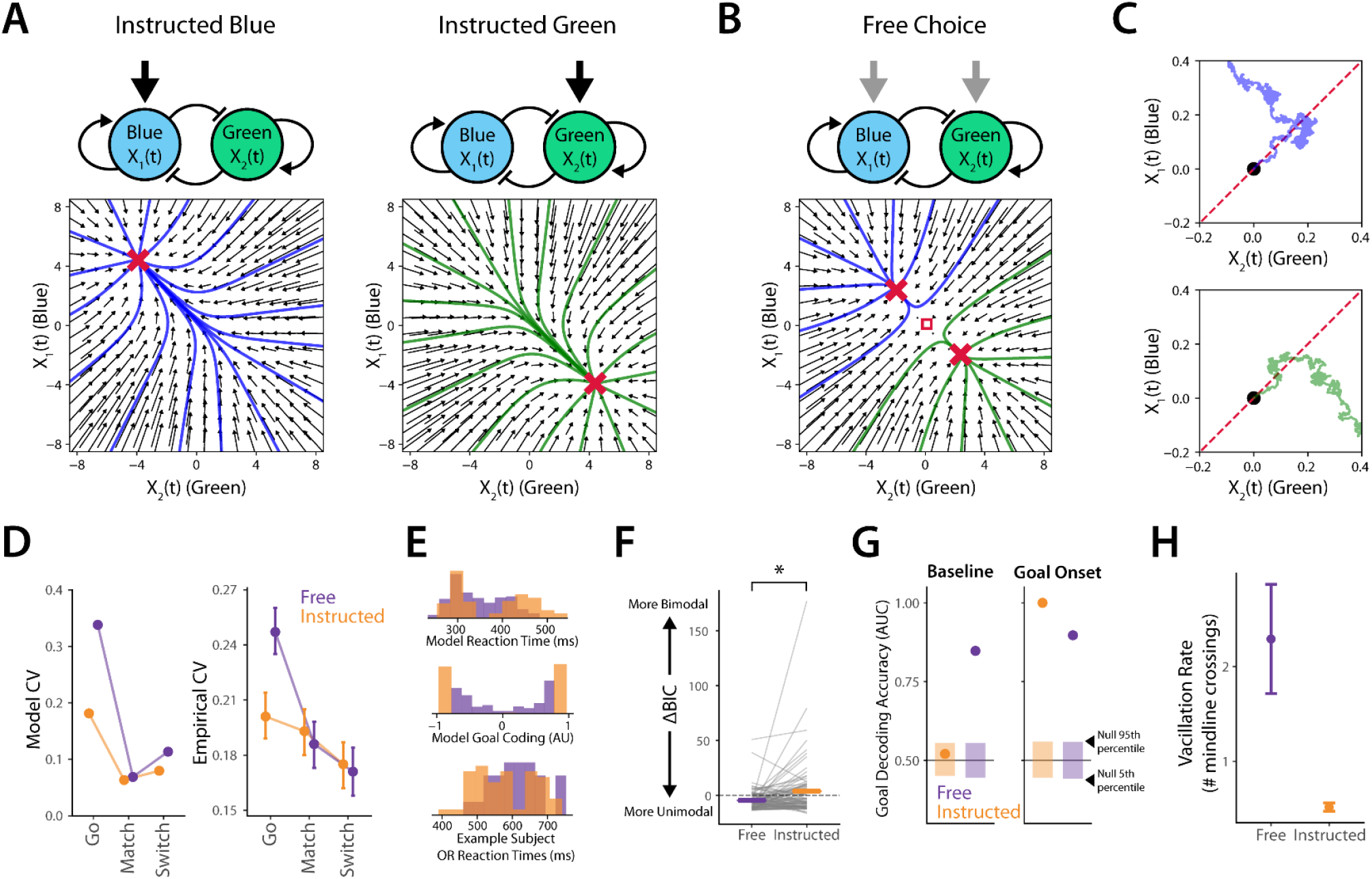
Symmetry-breaking winner-take-all dynamics explain free choice behavior. **(A-B)** Vector plots and fixed points for the goal circuit. Black arrows reflect local gradients as a function of node activities. Colored trajectories reflect simulated trajectories (without input noise) from a variety of initializations (Actual simulations for modeling behavior were initialized at the origin and had a small amount of input noise). Red Xs indicate fixed points. **(A)** Asymmetric input in Instructed conditions lead to single fixed points. **(B)** Symmetric input leads to the emergence of bistability in the winner-take-all circuit. The red square at (0,0) indicates the window for the visualization (or inset) in panel C. **(C)** Visualization of two example simulated Free choice trials with noise, zoomed into the red square in B. In both cases, the activity fluctuates around the initialized location (black circles at (0,0)) until noise drives the circuit towards one of the two attractors enough for the WTA dynamics to pull it into that basic of attraction. **(D-F)** Validating novel model predictions in empirical data. **(D)** Reaction time coefficient of variation (CV; standard deviation /mean) for model (left) and empirical (right) data. Empirical CVs were calculated per-subject and then averaged (mean & 95% confidence intervals from LME models). **(E)** Top: histogram of model RTs for all free (purple) and instructed (orange) override trials. Instructed trials were more bimodal than free. Middle: histogram of goal coding prior to Map onset (goal coding defined as the difference in activity between the green and blue goal nodes). Bottom: histogram of an example participant’s RTs on all override trials, with clearly stronger bimodality in instructed compared to free context. **(F)** The difference in Bayesian Information Criterion (BIC; model-fit metric) between unimodal and bimodal models. Positive (negative) ΔBIC indicates more evidence for bimodality (unimodality). **(G-H)** WTA dynamics can explain other free-choice phenomena. **(G)** Goal decoding performance at pre-goal onset baseline (model activity 500 ms before goal onset) and immediately following goal onset (model activity 100 ms after goal onset). Circles denote average decoding across 5 cross-validation folds. The semi-transparent bars mark the distribution of decoding accuracies obtained from a null distribution, where we trained decoders on activity with shuffled labels (1000 bootstraps; rectangles give from the 5^th^ to the 95^th^ percentile). **(H)** Vacillation rates in Free and Instructed Go trials, defined as the number of crossings of the midline/separatrix (dotted red line in panel C) between Goal onset and Response Map (mean & 95% confidence intervals across trials).

In Instructed trials, circuit inputs are asymmetric, leading to a single stable fixed point where the node that the network was instructed to select has high activity and the other has low activity (Fig. 3A, Fig. S3A). However, in Free trials, circuit inputs are symmetric, leading to the emergence of two stable fixed points, corresponding to the two possible goals (Fig 3B). Crucially, these two fixed points in the Free context are less extreme (in terms of the magnitude along the axes) than the fixed points in the Instructed context, meaning that freely chosen goals will be encoded less strongly than instructed ones even after commitment (Fig. S3B). This encoding difference percolates through the hierarchical circuits, leading to less activation in the conjunction and motor circuits and, thus, slower reaction times for Freely chosen compared to Instructed goals (Fig. S3A, B). This weaker encoding also leads to relatively lower goal-switching costs for Free compared to Instructed goals, because there is less activity in the conjunction circuit for an override signal to overcome on Free trials (Fig. S3A, B). Importantly, under this model, no matter how much time is given for participants to make their choices, freely selected goals will always result in longer reaction times and lower goal-switching costs compared to instructed goals.

How is input symmetry broken in Free Choice? The circuit is initialized at the origin (0,0), and near the origin symmetric input leads to very weak local gradients (Fig. 3B). Therefore, the small amount of noise we add to simulations plays a dominant role in determining the system’s trajectory: the system takes a random walk (with weak WTA dynamics) until it veers far enough from the line dividing the basins of attraction of the two attractors (dashed red line in Fig. 3C, known as the separatrix) for the attractor dynamics to overcome noise fluctuations; the trajectory then heads towards one of the two fixed points (Fig. 3C). In contrast, in Instructed contexts the input signals dominate and noise has a very minor role. Crucially, this process leads to a slower and more variable convergence to fixed points for Free compared to Instructed goals (Fig S3A, B).

Can symmetry-breaking attractor dynamics explain signatures of free choice in our behavioral data? Because the model was designed specifically to reproduce our core behavioral findings, it exhibits longer RTs and smaller Switch costs for Free contexts by design. To rigorously test the model’s validity, we examined its emergent properties—behavioral predictions that naturally arise from the network’s architecture but were not optimized for during fitting. We reasoned that because the goal-selection process in Free contexts is driven by stochastic noise (unlike the signal-dominated Instructed contexts), this mechanism should result in more variable RTs for Free than for Instructed contexts, even when considering the longer average RTs for Free compared to Instructed trials. We therefore tracked the coefficient of variation (CV; RT standard deviation /mean RT) in each condition. In simulated trials, Free Go trials had higher CV than all other conditions (Fig 3D left). In our empirical data, we also found that Free Go trials had the highest CVs among all conditions (Fig 3D right; p < 0.001; LME analysis post-hoc tests with Tukey correction), indicating that our model successfully predicted a novel signature of free choice behavior. Our model also predicted that Switch conditions would have slightly greater CVs than Match conditions (Fig. 3D left), but the empirical data did not bear out a clear pattern (Fig. 3D right). The empirical differences in CVs between Match and Switch conditions were generally unreliable (only the Instructed Match vs. Free Switch comparison reached significance, p = 0.035; for all other Match-Switch comparisons p > 0.05; LME with Tukey correction). This result suggests other mechanisms may influence RT variability on override trials.

We next tested whether our model could predict a novel difference in goal-switching behavior. We reasoned that because freely chosen goals converge more slowly and are encoded more weakly than instructed goals, they should be easier to override. Consequently, the RTs across all Free override trials, including those where participants were undecided, should blend into a single, unimodal RT distribution when pooled together. In contrast, the strong encoding of Instructed goals should result in fast Match RTs and slow Switch RTs, producing a distinctively bimodal distribution (Fig. 3E top). Importantly, our model shows that this unimodal blending in the Free context emerges downstream in the conjunction circuit, after response-map onset, because the goal circuit shows strong bimodal encoding prior to response-map onset (Fig. 3E center).

To test whether empirical RTs exhibited this emergent pattern, we quantified bimodality in our human data by fitting each single-participant override RT distributions with a Gaussian mixture model containing either one (unimodal) or two (bimodal) components (see an example participant’s results in Fig. 3E bottom; see Methods for details). Comparing the Bayesian Information Criterion (BIC) between these models confirmed our model’s prediction. We found significantly more evidence for bimodality in Instructed override trials compared to Free override trials (Fig. 3F; mean±SEM; Empirical Free ΔBIC = −4.52±1.15; Instructed ΔBIC = 3.55±2.9; W=1077, p=0.009, Wilcoxon sign-rank test; model Free ΔBIC = 16.95, Instructed ΔBIC = 192.05). This implies that symmetry-breaking attractor dynamics can predict not only basic RT differences, but also how dynamically Free and Instructed goals interact with shifting behavioral demands (i.e., the introduction of override signals).

Symmetry-breaking attractor dynamics also offer a parsimonious account of two signatures of free choice behavior that have been previously reported in the literature: pre-stimulus choice decoding^9,20^ and vacillation^33–35^. In our attractor network model, as in the brain, activity does not begin at goal onset but rather has pre-stimulus components (see the activity before 0 in all panels of Fig. S3 for example). For instructed trials, goal input is powerful enough to override such pre-stimulus activity. But for Free trials, pre-stimulus activity may determine the later decision: if noise has, by chance, driven the network closer to one of the fixed points prior to the goal input being received, it is more likely that the corresponding goal will be selected after the goal input is received, because the input is symmetric. Indeed, in our model, we were able to predict later goal selection based on the pre-stimulus simulated activity for Free but not Instructed Go trials (Fig 3G left; 5-fold cross-validated decoding AUC; p < 0.05 for Free, p > 0.05 for Instructed, permutation test; see also separation between activity at Goal Onset in Fig 2B top left and right).

As expected, 100 ms after goal onset we observed high goal decoding for both contexts (Fig 3G right; both p < 0.05, permutation test). Interestingly, we see that while the decoding in the Instructed condition is near perfect, the decoding in the Free condition is only around 90% accurate and not that much higher than in the baseline period. This is not surprising given the slower dynamics in Free compared to Instructed (compare the difference between blue and green activations 100 ms after goal onset in the Goal Circuit in Fig. S3A and B) and the locations of the fixed points in the two conditions (Fig. 3A vs. B), meaning that goal encoding is overall stronger for Instructed than Free contexts.

Beyond pre-stimulus decoding, symmetry-breaking attractor dynamics also account for vacillatory behavior during free choice. Individuals sometimes reconsider or go back-and-forth between choice options prior to committing when making free choices ^33–35^. We wondered whether our model could explain this phenomenon. In our model, symmetric input near the origin produces weak gradients, enabling noise to drive trajectories back and forth across the separatrix before attractor dynamics dominate. We quantified vacillation as the number of trajectory crossings of this midline and found it was significantly greater for Free, with an average of 2.29±0.29 midline crossings per trial (mean±SEM), compared to Instructed trials which had an average of 0.52±0.022 crossings per trial (Z=6.51, p<0.001; rank-sums test, Fig 3H), consistent with empirical reports. The nature of these vacillations—whether between an undecided state and a single goal, or between competing goals— remains an open question we address in the Discussion. Altogether, however, we find that symmetry-breaking attractor dynamics can successfully account for a range of novel and previously reported signatures of Free Choice behavior.

### Behavior Replicates in an Independent Cohort

To replicate our results and study the neural mechanisms underlying them, we recruited 30 additional participants who carried out our abstract free choice task, while we recorded electroencephalography (EEG) data. The task they performed was identical to the one described previously, with two exceptions designed to optimize for EEG decoding. First, to increase our trial count and statistical power, we omitted the early SOA periods and exclusively utilized SOAs uniformly distributed in the 700 to 1200 ms range. Second, to further improve decodability, the goal stimuli disappeared from the screen 400 ms after goal onset ^36^.

Participants exhibited the same general pattern of RTs as in our prior task (Fig 4A; compare to Fig. 1B). Free Go trials exhibited an average reaction time of 510.3 ms [470.7, 549.8], which was significantly slower than Instructed Go trials (mean = 455.9 ms [416.5, 495.3]; T(10887.1)=-17.95, p<0.001, LME post-hoc tests with Tukey correction). Likewise, Free Switch trials (mean = 564.1 ms [522.8, 605.4]) were significantly slower than Free Match trials (mean = 503.9 ms [463.4, 544.4]; T(10887.7)=6.98, p<0.001), and Instructed Switch trials (mean = 610.1 ms [570.1, 650.1]) were significantly slower than Instructed Match trials (mean = 481.8 ms [441.9, 521.7]; T(10887.3)=22.72, p<0.001). Replicating our core finding, switch costs were significantly larger for Instructed compared to Free contexts (126.9 ms vs 59.7 ms; W=11.0, p<0.001, Wilcoxon sign-rank test).

**Figure 4.**
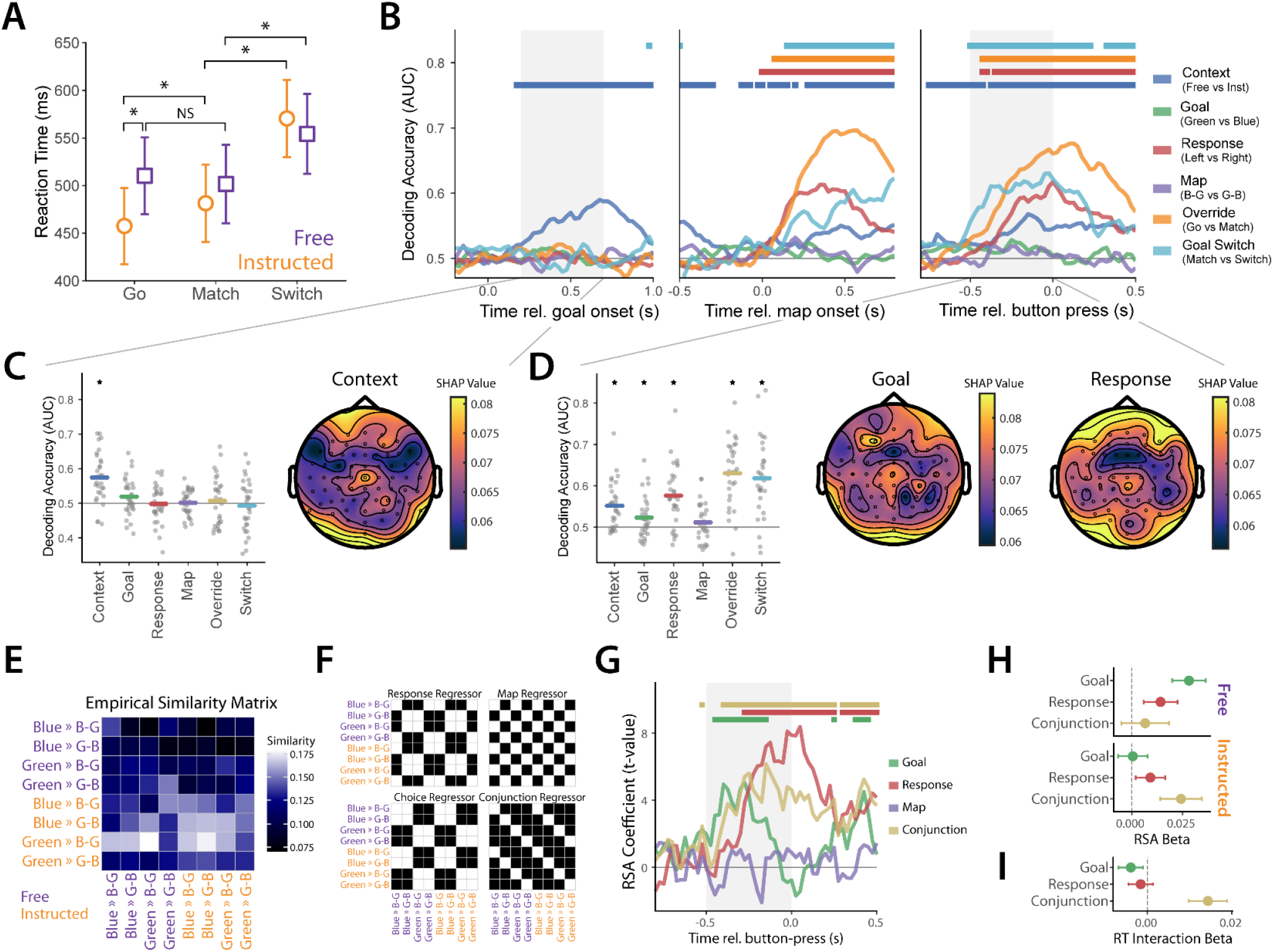
Conjunctive representations facilitate translation of abstract goals to motor output. **(A)** Mean & 95% confidence intervals for RTs in the EEG study (N=30; mean, CI, & significance assessed from LME analysis). All main behavioral features (difference between Free & Instructed RTs, greater switch costs for Instructed than Free contexts) replicated. **(B)** Temporal decoding analysis of context, goal, response, map, override, and goal shift relative to the time of goal onset (left), response-mapping onset (middle), and button press (right). Curves depict cross-subject average decoding accuracy, same-color horizontal bars above denote periods of significant decoding (regression against 50%, FDR corrected). Grey-shaded regions denote periods for further analysis in panels C and D. **(C)** Single-subject (grey dots) and average decoding accuracies (colored bars) for interval (0.2 s, 0.7 s) relative to goal onset (see B, left). Significance denoted with t-test against 50%. Right: Topoplot of average Context decoding SHAP values for each recording site (averaged across frequencies and then across participants). **(D)** Left: Decoding accuracies (as in C) in the interval (−0.5 s, 0 s) relative to button-press (see B, right). Right: SHAP Topoplots (as in C) for Goal (left) and Response (right) decoding. **(E)** Empirical decoder-based similarity matrix for representational similarity analysis (cross-validated confusion matrix averaged across participants). For this analysis we used correct Go trials only, and categorized the data according to the context, goal, and map, which also defined the given response (e.g. Blue >> B-G in orange indicates an instructed Blue goal trial that was followed by a B-G response map, and thus a leftward response). **(F)** RSA regressor matrices. White indicates value of 1, black indicates value of 0. Regressor matrices indicate what the empirical similarity matrix would be if the brain cared only about the given regressor (e.g. if the brain cared only about the Response, it would treat Blue >> G-B and Green >> B-G as equivalent, as they both entail rightward responses). The regressor matrices can be mapped onto the empirical similarity matrix to investigate which features are encoded in neural activity. **(G)** Time-resolved RSA. Trajectories are time-stamp-by-time-stamp RSA coefficients aligned to button press (higher values indicate stronger confidence of a match between the neural pattern and that variable’s corresponding regressor matrix, in F). Horizontal bars at the top of the panel are periods of significance (p < 0.05, LME analysis, FDR corrected). Goal encoding emerges first, followed by the Goal-Map conjunction, followed by Response. The grey-shaded region is the window used for analysis in panels H-I. **(H)** Context-specific RSA. Means & 95% CI (LME analysis) of RSA estimated effects conducted separately for Free & Instructed conditions. Free conditions have relatively greater goal representation, while instructed conditions have relatively greater Conjunction representation. **(I)** Conjunctive representations relate to behavior. Means & 95% CI (LME analysis) for regressor-reaction time interaction beta coefficient. Conjunctive representations exhibited significant interactions with behavior.

Finally, participants’ accuracy was lower for Switch than Match in both contexts (Free Match: 99.0%, Free Switch: 93.3%, Instructed Match: 99.5%, Instructed Switch: 86.3%; p < 0.001 both, logistic mixed effects analysis post-hoc tests with Tukey correction). Furthermore, Instructed Switch accuracy was significantly lower than Free Switch accuracy (p < 0.001; same tests as above, Fig S4). Our second study thus successfully replicates the core behavioral finding that freely chosen goals produce intact but diminished signatures of goal commitment relative to instructed goals.

### EEG Signals Encode Task Variables

We next investigated the extent to which the brain (via EEG) encoded the key variables required for performing our task, as per our model. Several event-related potentials differed between Free and Instructed conditions, including the P300 following goal and map onset, the LRP prior to response, and the ERN (Fig S5-8), indicating that neural representations common to decision-making and control tasks are also present in our dataset. To investigate how EEG signals encoded task variables in a signal-agnostic manner, we applied a cross-validated decoding procedure to time-varying (Fig 4B) and time-averaged (Fig 4C-D) frequency-specific power at all recorded electrode locations (see Methods). Decoding targets included Context (Free vs. Instructed), Goal (Blue vs. Green), Response (Left vs. Right), and Map (B-G vs G-B) on Go trials. We also targeted Override signals by contrasting Go and Match trials (which share the same goal, but lack or include an override cue, respectively), and Goal Switching by contrasting Match and Switch trials. Together, the latter two allowed us to separately target override-based control and goal switching.

During the delay period between Goal Onset and Response Map, we were able to decode only the Context (Fig 4B), reaching an average AUC of 0.575±0.013 (mean±SEM) across participants, which was significantly greater than chance level of 0.5 (Fig 4C; W=427.0, p<0.001, one sided Wilcoxon sign-rank test, all other targets had p>0.1 in this period). In the 500 milliseconds before button press, we were able to significantly decode Context (mean±SEM; AUC=0.552±0.010, W=424.0, p<0.001), Goal (AUC=0.523±0.009, W=332.0, p=0.020), Response (AUC=0.575±0.013, W=439.0, p<0.001), the presence of an Override signal (AUC=0.631±0.015, W=458.0, p<0.001), and Goal Switching (AUC=0.618±450.0, W=450.0, p<0.001). We could not significantly decode the response Map for this time range (AUC=0.511±0.008, W=286.0, p=0.140; Fig 4D).

Overall, these results indicate that the main signals needed to accomplish our task were encoded at the expected periods. Notably, however, participants’ abstract goal was not reliably decodable at individual timepoints (Fig. 4B), only in time-averaged activity in the 500 ms before response onset (Fig. 4D). This implies that Goal representations were encoded only subtly in EEG signals, which is consistent with prior work showing abstract choice decoding in the high 50%-low 60% range even with higher-SNR methods^13,37^.

We next considered what regions might be more involved in representing these different variables. For every decoding feature (electrode-frequency pair) and decoding target in the windows of interest (grey windows in Fig 4B) we extracted the SHAP value, a measure of the specific feature’s importance for decoding a target variable. We then averaged the absolute SHAP values across frequencies to investigate the spatial distribution of decoding-relevant information (Fig 4C-D; see Fig S9-10 for frequency-band specific plots). Context SHAP values were particularly high over fronto-central regions (electrode Cz), commensurate with prior decoding efforts for Free vs. Instructed choices ^15^. We next considered Goal and Response signals (Fig 4D). Although the broader spatial distributions of SHAP values were distinct for these two targets, both shared an overlapping peak of around Cz. We thus hypothesized that these fronto-central circuits are not only important for independently encoding Goals and Responses, but may also play a role in translating abstract Goals into actionable motor response plans—which we explicitly tested in the following section. While these standard decoding approaches confirmed the presence of isolated task variables, they cannot reveal how these variables are integrated.

### Conjunctive Representations Translate Abstract Goals to Motor Output

How are abstract goals translated into motor output? Our hierarchical attractor network model achieved this using a conjunctive circuit that integrates abstract goals with relevant sensory information (the response map) to trigger a specific action. We tested whether neural signals encoded features of such conjunctive representations during our task using a decoding-based representational-similarity analysis (RSA; see Methods), which has previously been used to characterize conjunctive representations during cognitive control ^28,29^. Briefly, we trained decoders to predict which of 8 classes a given Go trial belonged (2 contexts x 2 goals x 2 maps; note that the response on a correct trial is a direct combination of goal and map). From these confusion profiles we constructed a similarity matrix (Fig. 4E), which we regressed on exemplar matrices that encode which combinations of conditions should be (dis)similar based on which variables are encoded in neural activity (Fig 4F).

Using this RSA approach, at every timestamp we tested the corresponding empirical similarity matrix (computed independently for that specific moment, unlike the time-averaged summary shown in Fig. 4E) against the four response regressors (in Fig. 4F). We found that EEG signals reliably encoded conjunctive representations prior to response onset alongside abstract Goal and motor Response signals (Fig 4G; all p < 0.05, LME analysis). Importantly, encoding onsets were temporally ordered within the last 500 ms before button press, with Goal encoding emerging first, followed by Conjunction, and then Response—in line with the temporal order predicted by our model (Fig. S3). The Map was not significantly encoded at any point (p > 0.05).

Interestingly, when we conducted this analysis separately for the Free and Instructed conditions, we saw relatively greater Goal encoding for Free contexts and relatively more Conjunction encoding for Instructed contexts (Fig. 4H). At first glance, greater Goal decoding for Free trials might seem to contradict our model’s prediction that freely chosen goals converge on less extreme fixed points (i.e., weaker representations). However, because Free goals develop more slowly (Fig. S3A, B), the brain dwells in this goal-selection phase longer, rendering it more readily decodable over the pre-response window. Conversely, the stronger Conjunction encoding for Instructed trials directly aligns with our model’s predictions: deeply entrenched Instructed goals robustly drive the rapid activation of downstream sensorimotor conjunctive circuits.

To test whether the conjunctive representations played a role in translating abstract goals into motor output, we next investigated how encoding interacted with reaction time during the pre-response window. We found that conjunctive representations significantly interacted with RT during the pre-response period (β = 0.0143, p = 0.002; LME analysis), while Goal and Response representations did not (β = −0.0039 and β = −0.0016 respectively, both p > 0.1). This positive Conjunction×RT interaction indicates that longer reaction times are associated with stronger conjunctive encoding. Rather than implying that stronger encoding slows down responses, this relationship likely reflects a time-on-task effect: trials with longer RTs afford the brain more time to dwell in the conjunctive state, rendering the representation more stable and decodable. The temporal ordering of Goal, Conjunction, and Response representations alongside this specific relationship of conjunctive representations to behavioral variability provide evidence for the existence of something resembling a key element of our attractor network model in the brain—the conjunction circuit—and suggests that such conjunctive representations play a role in translating abstract goals into motor output.

## Discussion

The capacity to select between alternatives when there is no clearly superior option is paramount to everyday life. Such decisions must often be made abstractly, without an immediate translation of the decision alternatives into action plans. Nonetheless, indecision in such circumstances can be frustrating when trying to decide what to eat for dinner, or even dangerous when trying to avoid obstacles when driving. In the present study, we account for behavioral signatures of free choice via symmetry-breaking attractor dynamics, providing a circuit-level mechanism for the formation of free choices. More specifically, we developed a hierarchical attractor network model with this mechanism that accounts for the free choice behavior exhibited by participants in our abstract decision-making paradigm. We demonstrate that our model can reproduce key behavioral signatures of free choice; namely, longer RTs^1–6^ and lower goal-switching costs^22^ for free than instructed goals. The model also made a novel prediction —increased RT variability for free choice—which we successfully validated in the empirical data. It further parsimoniously explained pre-stimulus choice decoding and vacillation in free-choice decisions, which have been demonstrated in the literature. Finally, EEG analysis showed that key components of this hierarchical model are represented in the human brain.

Prior work investigating free choice attributed its longer response times to the added requirement of making a decision that is absent when just following instructions^1,5^. Contrary to this hypothesis, we found that participants exhibit slower RTs for freely selected goals even when goal selection and response mapping are separated by a considerable delay (up to 1.2 s). Our model attributes this difference to less extreme fixed points for free choice in a WTA circuit ^24,31,38^, given symmetric (in Free) compared to asymmetric (in Instructed) model inputs, which percolate downstream to slower motor activation. The convergence of model activity to fixed points is slower and more variable for free choice, which also explains why we observed similar but slower goal commitment (i.e., convergence to a fixed point) in Free compared to Instructed trials. Our model also suggests that above-chance choice decoding before options onset ^9,20^ reflects pre-stimulus attractor dynamics that influence the later decision under symmetric but not asymmetric input. Similarly, vacillation between response options after they are given in free choice ^33–35^ is due to noise-dominated trajectories near the origin, where symmetric input produces weak gradients, and the system wanders before committing to a goal. Our model thus offers a parsimonious explanation for a range of behavioral and neural signatures of free choice.

We showed that key model components are encoded in human neural activity (via EEG). We were able to reliably decode Context (free vs. instructed), Goal (blue vs. green), and Response (left vs. right) directly from time-frequency EEG activity. Notably, goal decoding was low, indicating EEG does not have sufficient signal-to-noise ratio to fully characterize these representations. Nevertheless, studies using methods with increased signal-to-noise ratio, such as fMRI or MEG, achieve higher accuracies— in the high 50% to low 60% range—for abstract choice ^13,37^. This implies that coarse, population-level, cortical-based signals do not provide sufficient information to fully characterize abstract choices. It is thus possible that invasive recordings might be required for a full characterization of the neural mechanisms underlying abstract decision-making.

Despite modest univariate decoding accuracies, we were able to establish the presence of different types of representations using representational similarity analysis (RSA). We found that human EEG signals encode conjunctive representations that, our model suggests, translate abstract goals into actions. Such conjunctive representations emerge when information regarding multiple variables must be integrated to produce a response ^27–29^ and are related to event file theories of action. For instance, Kikumoto and colleagues ^28^ found that conjunctive representations emerged between and bridged stimulus and response representations. In our study, goal and response representations were similarly sequentially bridged by a conjunctive representation, and it was specifically this conjunctive representation that related to behavioral variability. Thus, in addition to facilitating stimulus-response mapping, our study suggests that conjunctive representations also enable the translation of abstract goals to motor output.

We did not find evidence for specific response-map representations, so how could such conjunctive representations form? A trivial possibility is that the neural representations for response-mapping are inaccessible using EEG. A more interesting possibility is that participants form a contingency representation for how to respond: Ehrlich and Murray ^39^ showed that participants and artificial neural networks pre-compute a decision about how to respond to an upcoming stimulus given prior information rather than waiting to make decisions later. For instance, given a task requiring comparison of two sequential stimuli, one might pre-plan how to respond to the second stimulus (e.g., in our case, for a Blue goal the participant would plan to respond left /right for a B-G /G-B map, respectively) rather than waiting for the mapping to be revealed to plan a response. Similarly, our model supposes that the abstract goal activates compatible conjunctive nodes, effectively forming a contingency representation to respond left for one layout and right for the other. However, in our experimental design, a specific abstract goal is perfectly correlated with a specific contingency rule. For example, every time a participant commits to the abstract goal of “Blue,” they must simultaneously adopt the exact same contingency rule (“press left for B-G, right for G-B”). Because these two cognitive states always cooccur and are completely confounded, our current neural decoding cannot disentangle the representation of the abstract goal from the representation of the pre-planned rule. Investigating the precise relationship between goal, conjunctive, and contingency representations will therefore require future tasks designed to separate these variables.

To explain why Match trials are slower than Go trials in the Instructed context, our model required a direct inhibitory pathway from the override to the response circuits. This mechanism allows for the rapid but transient suppression of motor output the moment an override cue appears. Biologically, this is reminiscent of the “hyperdirect” pathway—a cortico-subthalamic circuit known to trigger rapid action cancellation ^25,26^. However, recent work by Kikumoto and colleagues ^40^ suggests a competing mechanism: they found that action stopping suppresses intermediate conjunctive representations rather than final motor response representations. Our model reconciles these findings by suggesting both mechanisms operate in tandem. Specifically, the conjunctive suppression Kikumoto and colleagues observed corresponds to how our model’s conjunction circuit is actively disrupted during Switch trials, whereas the hyperdirect pathway handles the transient, raw motor inhibition seen in Match trials. Future research combining goal-switching paradigms with, again, intracranial recordings could definitively test whether inhibitory control operates on intermediate conjunctive representations, terminal motor representations, or a combination of both.

Other phenomena associated with free choice may prove a challenge to our model. First, although our model can explain internal vacillation, participants making free choices sometimes go further: they make overt movements and then fully inhibit them ^34^. Given the stability of fixed points in an attractor network, it is not clear how our model would account for such overt behavioral switching once a decision state is reached. Second, people learn more from freely chosen events than from following instructions ^41,42^, a phenomenon thought to be driven by stronger reward-prediction errors for free compared to instructed choices ^42^. Our model suggests that freely chosen goals are more weakly encoded than instructed ones. Thus, if reward prediction errors are derived from the mismatch between expected and actual results, our model would incorrectly predict smaller reward prediction errors for free choices. Therefore, a distinct mechanism likely underlies these learning differences. Finally, freely chosen actions are often associated with greater sense of agency than instructed actions ^43^. If agency judgments similarly rely on a mismatch computation ^44^, our model’s weaker fixed points would again incorrectly predict lower agency for free choices. Nevertheless, the breadth of phenomena that our model successfully and parsimoniously accounts for suggests that symmetry-breaking attractor dynamics play an important role in free choice. How these additional phenomena relate to the circuit mechanisms we identified is an important direction for future research.

Our attractor model offers a computational framework for investigating free choice. Conversely, free choice paradigms offer a useful method for studying attractor dynamics in human neuroscience. Attractor networks are a powerful and well-established tool for explaining human cognition ^45,46^. They offer parsimonious accounts for a range of cognitive phenomena, including working memory ^47–49^, perceptual and value-based decision-making ^23,24,50–52^, and other phenomena ^22,53,54^. Winner-take-all circuits have been extensively utilized as a model of perceptual and value-based decision-making^9,38^, and our work extends the utility of these models to account for abstract free choice phenomena. Our model suggests that the variability associated with symmetric input is a feature, not a “bug”, of winner-take-all circuits. Crucially, deviations in the activity of such attractor networks are implicated in psychiatric illnesses such as schizophrenia ^55,56^. Schizophrenia is also associated with impaired behaviors that may be related to free choice mechanisms, including random exploration ^57^ and the balance between model-based and model-free decision-making ^58^. Thus, free choice paradigms may be particularly useful for studying the intrinsic dynamics of attractor networks in the brain and how they relate to psychiatric illness.

## Data Availability

The data that support the findings of this study will be made available upon publication.

## Code Availability

The code that supports the findings of this study will be made available upon publication.

## Acknowledgments

We thank Lexi van der Hoeven for assistance with experiment design. We thank Emma Chen for assistance with data collection and EEG data preprocessing for Study 2. We thank Tomas Dominik, Lucas Jeay-Bizot, and Anne Loffler for valuable discussion of these results and their interpretation. J.G., A.S., and U.M. were supported by the John Templeton Foundation and the Fetzer Institute (Consciousness and Free Will: A Joint Neuroscientific-Philosophical Investigation (John Templeton Foundation #61283; Fetzer Institute, Fetzer Memorial Trust #4189)). J.G. was supported by the NSF (BCS-2219800) and fellowships from Chapman University and Cedars-Sinai Medical Center.

## Author Contributions

J.G., A.S., and U.M. conceptualized the project and designed the experiment. T.L. wrote the experiment scripts and oversaw data collection for Study 1. J.G. oversaw data collection for Study 2, performed data analysis for Study 1 & 2, designed the computational model and conducted model simulations and analysis, and wrote the initial manuscript draft. A.S. and U.M. supervised and provided feedback on data analysis. All authors edited the manuscript before submission.

## Methods

### Participants

For study 1, we recruited 80 participants (mean age 27.2 years, range: 18-80; 51.2% female, 47.5% male, 1.3% other) online through Prolific. These participants were compensated monetarily for their participation. For study 2, we recruited 31 participants (mean age 19.0 years, range: 18-23; 90% female, 10% male) through the Chapman University psychology research pool. These participants received course credit for their participation. One participant was excluded due low EEG signal quality. All participants provided informed consent. The experimental protocol was approved by the Chapman University ethics committee (IRB 20-115).

### Paradigm

Study 1: We ran a variant of a delayed-match-to-sample task, where participants were given a goal cue, waited a variable amount of time (see Response Map delay below), and then had to respond to the Response Map depending on the goal cue they were given (Figure 1). Our experiment could be described as a 2 × 2 × 3 design (Context [Free, Instructed] × delay [regular, early] × Map Type [Go, Match, Switch]). Trials were shuffled-counterbalanced to ensure consistent numbers of each trial type for each participant. The paradigm was implemented using Psychopy’s online toolbox.

Context: There were two types of goal cues, both consisting of colored discs—blue or green—placed above and below a fixation cross. 2/3 of all trials were Instructed Context, in which both discs were the same color, indicating that participants should press in the direction consistent with that color when the Response Map appeared. Hence, if the goal cue depicted two blue discs, participants should press left or right based on whether the blue disc was on the left or right side of the Response Map, respectively. The remaining 1/3 of trials were Free Context, in which the two discs were of different colors (one blue, one green), indicating that participants were free to select their own Goal (blue or green) and pursue it when the Response Map appeared.

Response Map delay: In this condition the participants were instructed to decide immediately on the color. There was a variable delay between the goal cue and Response Map. In regular trials, the delay was uniformly distributed between 700 and 1200 ms (70% of trials). After the first half of trials had been completed and participants habituated to the timing of the Maps, we introduced early delays of 200, 300, 400, or 500 ms on a subset of trials (each equally probable, 30% of trials overall). The goal cue remained onscreen until the Response Map was presented. The first 40 participants were run without the 200 ms delay, which was added after we determined that shorter delays would be informative.

Map Type: The Response Map again consisted of one blue and one green disc placed to the left and right of the fixation cross (B-G or G-B layout; each with 50% probability). Participants were instructed to press in the direction of the Instructed goal color or their freely chosen Goal. On Go trials (70%), both discs had a + sign overlaid, indicating participants should proceed with their prior Goal. On 30% of trials, an X appeared over one disc (Override trials), indicating that participants should press in that direction regardless of their prior Goal. Override trials were classified as Match trials when the cued direction was consistent with the participant’s prior Goal, and Switch trials when inconsistent. For Free trials, participants reported their intended Goal after each Override trial on a 5-point scale (1 = Decided Blue, 2 = Leaning Blue, 3 = Undecided, 4 = Leaning Green, 5 = Decided Green). We refer to Go, Match, and Switch collectively as Map Type.

Study 2: The paradigm was identical to Study 1 with the following exceptions: we omitted the early delay period and used only regular delays (700–1200 ms) to increase trial counts for EEG decoding, and goal cue stimuli disappeared from the screen 400 ms after onset to improve decodability of maintained goal representations (Chota et al., 2023).

### Procedure

Participants first completed 5 practice Instructed trials, then 5 practice Free trials, then 5 practice Override trials (both Free and Instructed), then 10 intermixed practice trials before beginning the main experiment of 8 blocks of 50 trials. Early delays were introduced after four blocks had been completed in Study 1.

### Outlier Removal and Data Retention

For each participant, within each delay bin (200, 300, 400, 500, 700–1200 ms) we removed trials with RTs more than 1.5 times the interquartile range from the median, that were incorrect (unless otherwise specified), or wherein participants reporting being “unsure” about their freely chosen goal following an override cue. For Study 1 we were left with an average of 347.7±36.5 trials per participant out of a total of 400 (mean ± stdev, range: 233-387). Of these, 47.5±2.5% of trials were Instructed Go trials, 10.6±1.5% were Instructed Match, 8.2±2.0% were Instructed Switch, 24.7±2.4% were Free Go trials, 5.3±1.7% were Free Match, and 3.7±1.7% were Free Switch. For study 2 we were left with an average of 364.1±19.7 trials per participant out of a total of 400 (mean ± stdev, range: 309-383). Of these, 47.9±1.7% of trials were Instructed Go trials, 10.0±1.2% were Instructed Match, 8.8±1.7% were Instructed Switch, 24.8±1.9% were Free Go trials, 5.3±1.8% were Free Match, and 3.2±1.6% were Free Switch.

### Mixed-Effects Analysis of Reaction Times

We analyzed single-trial data using linear mixed-effects models (LMEs) implemented using the Pymer4 package in Python, a port of the lme4 package in R. Post-hoc testing and F statistics used the package’s built-in functions with Tukey or Bonferroni corrections as indicated. For RT analyses during the regular delay period (Figure 1B) we used linear mixed effects regression with random intercepts per subject (RT ~ Context + MapType + Context * MapType + (1|subject)). For accuracy analyses on Override trials (Figure 1D) we used logistic mixed-effects regression with random intercepts (iscorrect ~ Context + MapType + Context*MapType + (1|subject)). For SOA dynamics analyses we used LMEs with random intercepts fit to a specific delay period (200, 300, 400, 500, 700-950, 950-1200) (RT ~ Context * MapType + (1|subject)). To estimate goal-switching costs at specific SOA bins (Fig 1G) we restricted our data to only Match and Switch trials in one context and fit LMEs with random intercepts (RT ~ MapType + (1|subject)). To estimate the SOA-specific difference between Free and Instructed RTs we fit LMEs with random intercepts to only Go trials at each bin (RT ~ Context + (1|subject)).

To estimate participant-specific goal-switching costs and their correlation with average Go RT (Figure 1C), we extracted random effects from a model with a random slope for MapType (RT ~ MapType + (MapType|subject)) on Override trials and then extracted random intercepts from a model fit only on Go trials (RT ~ (1|subject)). We performed this analysis separately for each context. Participant-specific coefficients (SwitchCost and AverageRT) were then correlated using standard regression.

### Hierarchical Attractor Network Model

We implemented a continuous rate model with 13 units governed by:

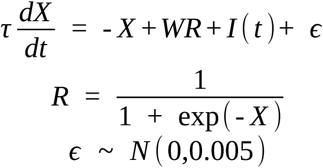

Where τ = 100 ms is the synaptic time decay constant, X ∈ ℝ^13^ is the synaptic current vector, R ∈ ℝ^13^ is the firing rate vector, W ∈ ℝ^13×13^ is the connectivity matrix implementing the circuit in Figure 2A, and I(t) ∈ ℝ^>13^ is the time-varying input encoding Context, Response Map layout, and Override direction.

The network comprised a Goal Circuit (Blue, Green nodes), a Map Circuit (B-G, G-B nodes), an Override Circuit (Left, Right nodes), a Conjunction Circuit (Blue-Left, Green-Left, Blue-Right, Green-Right nodes), a Response Circuit (Left, Right nodes), and a Response Inhibition node. Connectivity parameters: top-down excitatory connections (Goal → Conjunction = 1.26, Map → Conjunction = 1.27, Override → Conjunction = 1.17, Conjunction → Response = 6.83, Override → Inhibition = 76.34), lateral inhibition (Goal = −3.98, Map = −3.11, Conjunction = −2.09, Response = −1.81, Override = −1.85), self-excitation (Goal = 1.50, Map = 1.47, Conjunction = 0.53, Response = 0.79, Override = 0.63), global inhibition (Inhibition → Response = −16.52, Inhibition self = −20.52).

We simulated 1000 trials beginning with a 500 ms fixation period. For Instructed trials (50%), the cued Goal node received input of 1. For Free trials (50%), both Goal nodes received input of 0.5. After a delay of 700–1500 ms, Goal input ended and Response Map input began (value = 1). Override input (value = 1) was delivered on 50% of trials. Response was defined as a Response Circuit node exceeding R > 0.65; RT was measured from Response Map onset. Free choice Goals were labeled by the Goal node with higher firing rate at Response Map onset (difference < 0.05 labeled undecided; 3.56% excluded).

Model parameters were initially hand-tuned to generally reproduce behavior and then refined using simulated annealing and a composite loss function that penalized (1) the difference between model and empirical RTs (z-scored across conditions), (2) accuracy below 100%, (3) responses before Map onset, and (4) number of trials without either motor node crossing the response threshold. We simulated 500 iterations of 1000 trials each across 50 initial conditions, adding white noise to parameters at each iteration, and then selected the model that minimized the loss function after annealing. Across all iterations loss decreased from 3.46 to 0.52.

To plot state-space trajectories (Fig 2B), we defined orthogonal Goal and Motor axes by first computing the mean difference between average Blue and Green Go trial activity across all timepoints and the difference between Left and Right Go trial activity across all timepoints. The first difference constituted our Goal axis, and the component of the second difference that was orthogonal to the Goal axis defined our Motor axis. We computed explained variance as the variance of activity projected onto these axes divided by the overall variance of activity across time.

### Fixed-Point and Vector Field Analysis

We conducted fixed-point and vector field analyses of the Goal Circuit to characterize attractor dynamics under Instructed and Free Context conditions (Figure 3A–B). Stable fixed points were identified as locations where gradients equaled zero and the Jacobian had negative real eigenvalues. Vector fields were computed by evaluating dX/dt across a grid of initial conditions. Colored trajectories show simulated dynamics without input noise to illustrate convergence behavior.

### Coefficient of Variation Analysis

We calculated the coefficient of variation for the model as RT standard deviation divided by RT mean for each condition. We repeated this analysis for each subject and then fit LME to CV data (CV ~ Context + MapType + Context*MapType + (1|subject)) to get mean & confidence intervals (Fig 3D).

### Bimodality Analysis

To assess bimodality, we fit gaussian mixture models to log-RT distributions using the sci-kit learn package’s GaussianMixture command, with either 1 or 2 components for uni- or bi-modality respectively. We extracted the Bayesian information criterion from these fitted models and took the difference to get a single ΔBIC measure. We compared this measure across groups using a sign-rank test.

### Model Decoding Analysis

We used 5-fold cross-validated linear discriminant analysis (implemented through sci-kit learn) to assess whether the eventually chosen goal could be decoded before initial goals onset on Go trials. We then repeated this procedure 1000 times on data with shuffled labels to construct a null distribution (the rectangles in Fig 3G are between the 5^th^ and 95^th^ percentile of this distribution). For the Baseline period we used simulated activity from simulation onset to Goal onset (500 ms). For Goal Onset period we used simulated activity in the 100 ms after Goal onset.

### Vacillation Analysis

We defined the vacillation rate as the number of midline crossings in the goal circuit, i.e. the number of events where the sign of the difference between goal nodes (Blue – Green) changed from positive to negative or vice versa. We calculated the mean vacillation rate and 95% confidence interval for all Free and Instructed Go trials between Goal Onset and the Response Map (Fig 3H).

### EEG Acquisition & Preprocessing

EEG was recorded from participants in Study 2 using a 64-channel Biosemi system at an online sampling rate of 2048 Hz. The reference electrode was placed at Cz. Impedances were kept below 15 kΩ. Raw EEG data were preprocessed using Fieldtrip. Data were downsampled to 200 Hz and band-pass filtered (0.1-40 Hz). Continuous data were epoched 4 seconds around goal onset, response map onset, and button-press so that edge artifacts due to filtering could be omitted. We manually removed epochs with visible artifacts. Independent component analysis (ICA) was applied to remove blink artifacts.

### Time-Frequency Decomposition

Time-frequency representations (TFR) were computed using Morlet wavelets at 16 logarithmically spaced intervals from 2 to 40 Hz. We increased the number of cycles from 3 to 10 logarithmically as frequency increased. Power was taken as the squared absolute value of the resulting complex time series. For each trial and timepoint, power values across all electrode-frequency pairs served as features for decoding and RSA analyses.

### Decoding Analysis

We applied a cross-validated decoding procedure to classify task variables from single-TFR features. For time-resolved decoding (Fig 4B), a linear support vector classifier (SVC; kernel=‘linear’, class_weight=‘balanced’) was trained and tested using 5-fold cross-validation, and decoding accuracy was quantified as area under the ROC curve (AUC). Statistical significance at each timepoint was assessed using one-sided sign-rank tests against chance (AUC = 0.5), with FDR correction across time. For window-averaged decoding (Fig 4C–D), features were averaged across the specified time window before classification. Significance was assessed using one-sample t-tests against 0.5 across participants.

We decoded: Context (Free vs. Instructed), Goal (Blue vs. Green), Response (Left vs. Right), Response Map layout (B-G vs. G-B), Override presence (Go vs. Match), and Goal Shift (Match vs. Switch). Override and Goal Shift decoders were trained on Go+Match and Match+Switch trials respectively to avoid conflating variables.

To characterize spatial distributions of informative features, we computed SHAP (SHapley Additive exPlanations) values using a logistic regression classifier fitted to the full window-averaged dataset. SHAP values were averaged in absolute value across trials and then across frequencies to produce electrode-level importance maps (Fig 4C–D; Fig S9-10 for frequency-band-specific maps).

### Representational Similarity Analysis

We applied a decoder-based representational similarity analysis (dRSA) following Kikumoto et al. (2020; 2024). This approach constructs an empirical similarity matrix from classifier confusion profiles rather than raw pattern distances, enabling detection of both similarity and conjunctive representational structure.

For each participant, we defined 8 conditions corresponding to all combinations of Context (Free, Instructed), Goal (Blue, Green), and Response Map layout (B-G, G-B). We trained a Naïve Bayes classifier (implemented with sci-kit learn) to predict the probability a given trial belongs to each class using leave-one-out cross-validation at each timepoint. The resulting confusion matrix formed the empirical similarity matrix (Fig 4E).

We regressed the empirical similarity matrix against four regressor matrices encoding which condition pairs share the same Goal (Goal regressor), same Map layout (Map regressor), same combination of Goal and Map (Conjunction regressor), and same Response direction (Response regressor; Fig 4F). Because Conjunction on correct trials is a combination of Goal and Map, and Response is determined by the Goal and Map on correct trials, models would not fit all four factors simultaneously. Thus, we used LME models to examine how well single regressor matrices mapped onto the empirical similarity matrix (pred_prob ~ regressor + (1|subject)). Results were similar for the fitted factors when we fit multiple simultaneously. Significance at each timepoint was assessed using the fitted LME t-statistics with FDR correction across time. The grey-shaded window in Fig 4G (−500 to 0 ms relative to button press) was used for window-averaged RSA analyses in Fig 4H–I. For context-specific RSA (Fig 4H), we restricted the empirical similarity matrix to rows and columns belonging to the same Context (Free or Instructed), analyzing each 4×4 submatrix separately. For the RT interaction analysis (Fig 4I), we tested how different regressors interacted with within-subject z-scored reaction times (pred_prob ~ regressor * RT_z + (1|subject)). Same-condition pairs (i.e. the diagonal) were excluded for this analysis as their probability values tended to reflect stable condition discriminability rather than trial-level representational variation and therefore washed out behavioral effects. A significant positive interaction indicates stronger encoding is associated with longer RTs; a negative interaction indicates shorter RTs.

