## Supplementary Figures for "A neural circuit mechanism for abstract free choice"

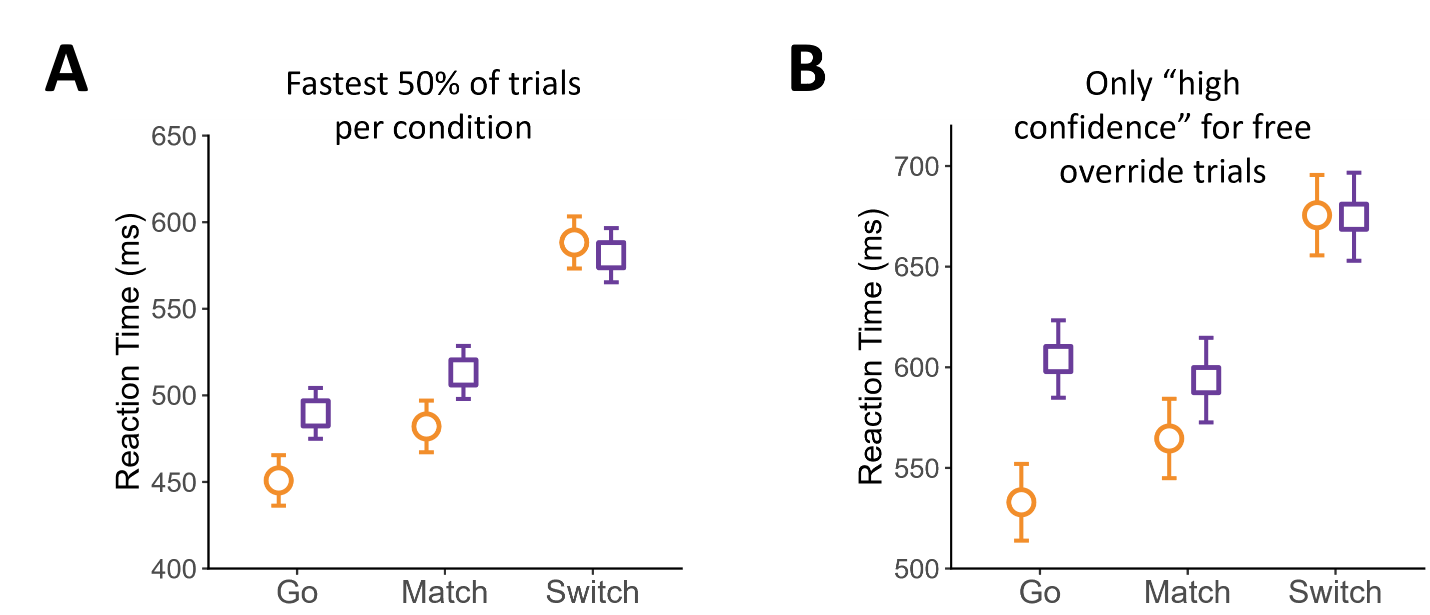

**Supplementary Figure 1.** Estimated condition mean RT & 95% confidence intervals (LME analysis) for control analyses. **(A)** We repeated our analysis from Fig 1B, retaining only the fastest 50% of trials for each condition for each participant. RT differences between Free (purple) and Instructed (orange) conditions were diminished but still present. Switch costs for free choice remained. This indicates that the observed effects were not due to long-tailed distributions in reaction times. **(B)** We repeated our analysis from Figure 1B, this time retaining only Free Override trials where participants indicated that they were sure they had chosen Green or Blue (as opposed to leaning towards one option). The pattern of results was again the same as in Fig. 1B.

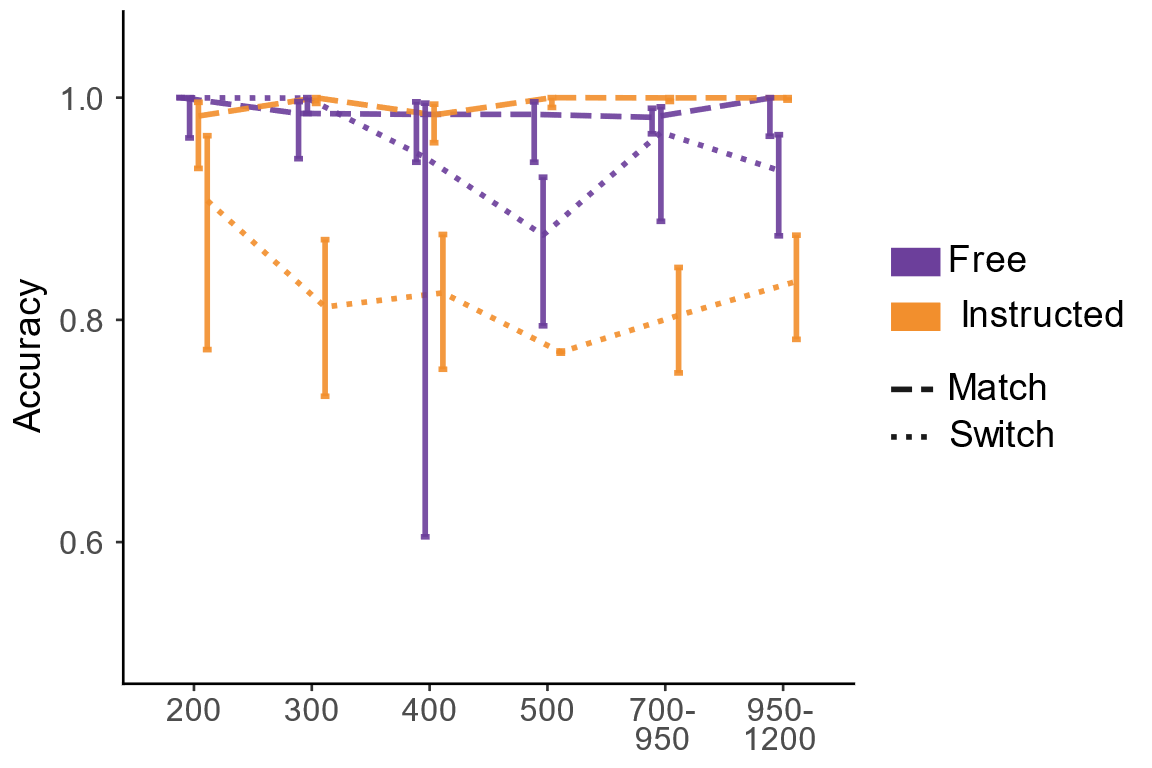

**Supplementary Figure 2.** Accuracy as a function of SOA in Switch trials (ratio of trials where participants were able to comply with the override cue that conflicted with the motor mapping for their original color choice). Average accuracy was calculated using logistic mixed-effects modeling (mean probability & 95% confidence intervals). Accuracy was high for all Match trials (dashed lines) and was relatively high at short SOAs for both Instructed and Free Switch trials (dotted lines). Instructed Switch accuracy fell to about 80% at SOAs of 300 ms, whereas Free Switch accuracy began dropping below ceiling at SOAs of 400 ms, indicating a difference in the dynamics of decision commitment between contexts.

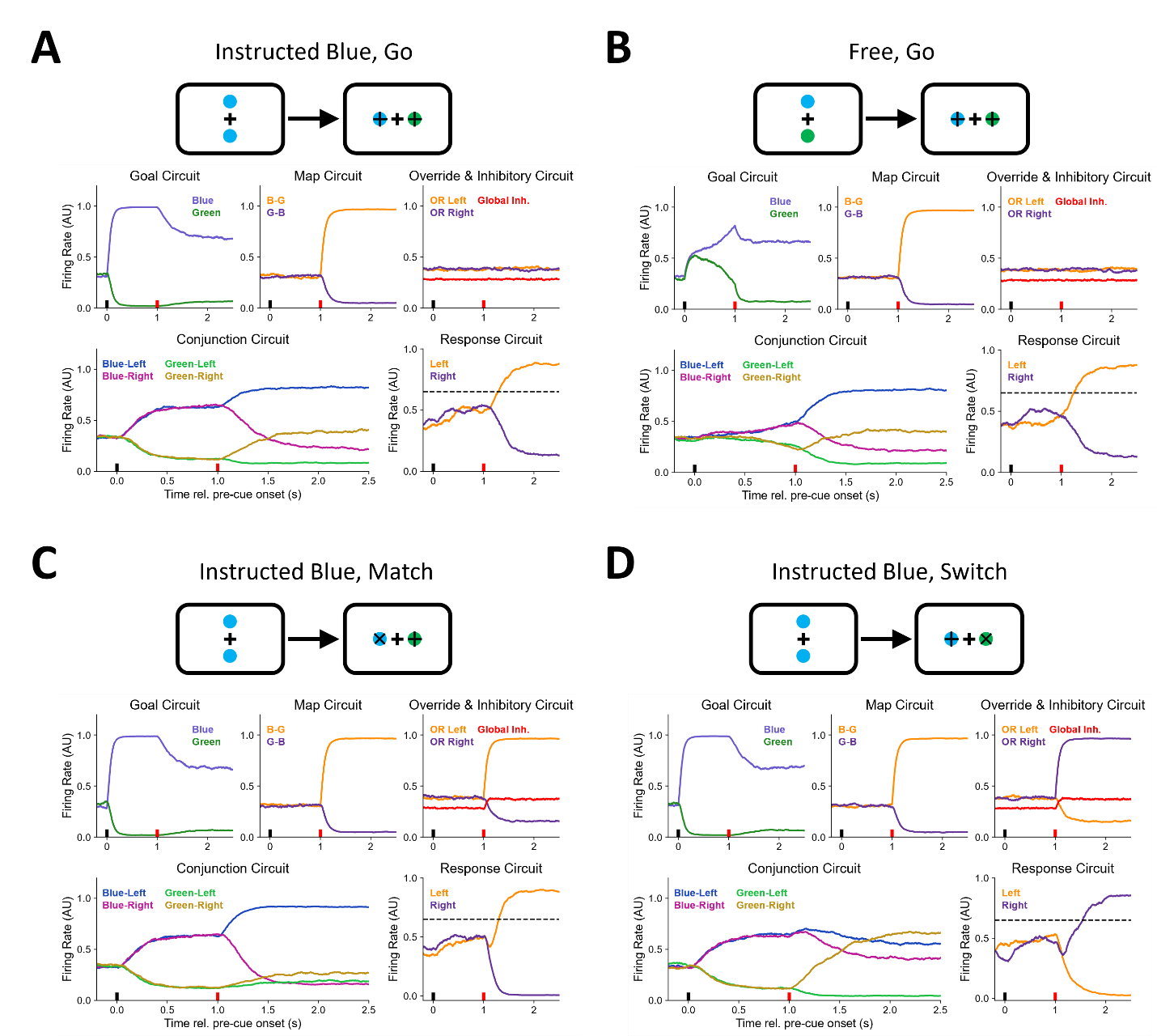

**Supplementary Figure 3.** Examples of simulated trials depicting the activity in all nodes (the Response Inhibition (Global Inh) activity is depicted in the Override & Inhibitory Circuit). In all cases, the short black line at time 0 indicates color-stimulus onset, and the similar red line at 1 is map onset (color-to-map duration varied in actual simulation but is constant here for visualization purposes). **(A)** Instructed Go trial, Blue goal. Per our circuit model diagram (Fig 2A), the selected goal (top left) innervates goal-consistent nodes in the conjunction circuit (bottom left). Then, upon activating the map circuit (top middle; blue is leftwards), the Conjunction circuit transitions into favoring the blue-left motor response, which leads the left motor node to cross the threshold (bottom right). The Override & Inhibition Circuit remains inactive (top left). **(B)** Free Go trial, Blue selected. A similar process takes place on Free trials, except for the slower convergence to a weaker fixed point, which leads to shallower activations in the conjunction circuit before map onset, and slower RTs following map onset. The Override & Inhibition Circuit again remains inactive. **(C)** Instructed Match trial. At map onset, an override cue is also applied (top right) which inhibits (in red) the Response Circuit through the global inhibitory pathway (see the small dip in the orange curve in the Response Circuit, bottom right), but largely leaves activity in the conjunction circuit intact, leading to slightly slower RTs. **(D)** Instructed Switch trial. In contrast to Match trials, override cues on switch trials disrupt the Conjunction Circuit representations (bottom left), in this case making the Green-Right activation eventually overtake the Blue-Left one. The network thus takes time to evolve and reach a supra-threshold activity in the Response Circuit.

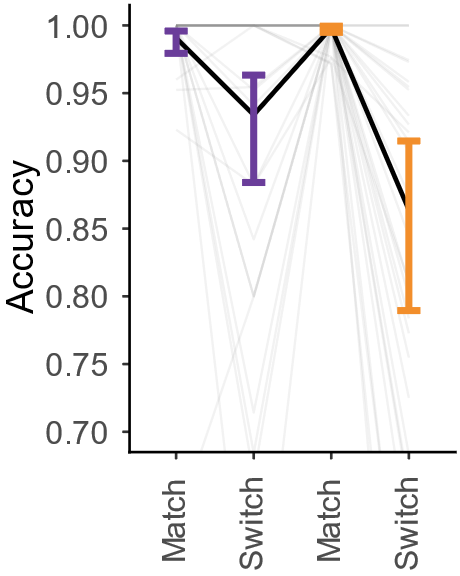

**Supplementary Figure 4.** Response accuracy on override trials for EEG study participants. Plotted and analyzed as in Figure 1D, means & 95% confidence intervals (estimated with logistic mixed effects; gray lines = individual participants).

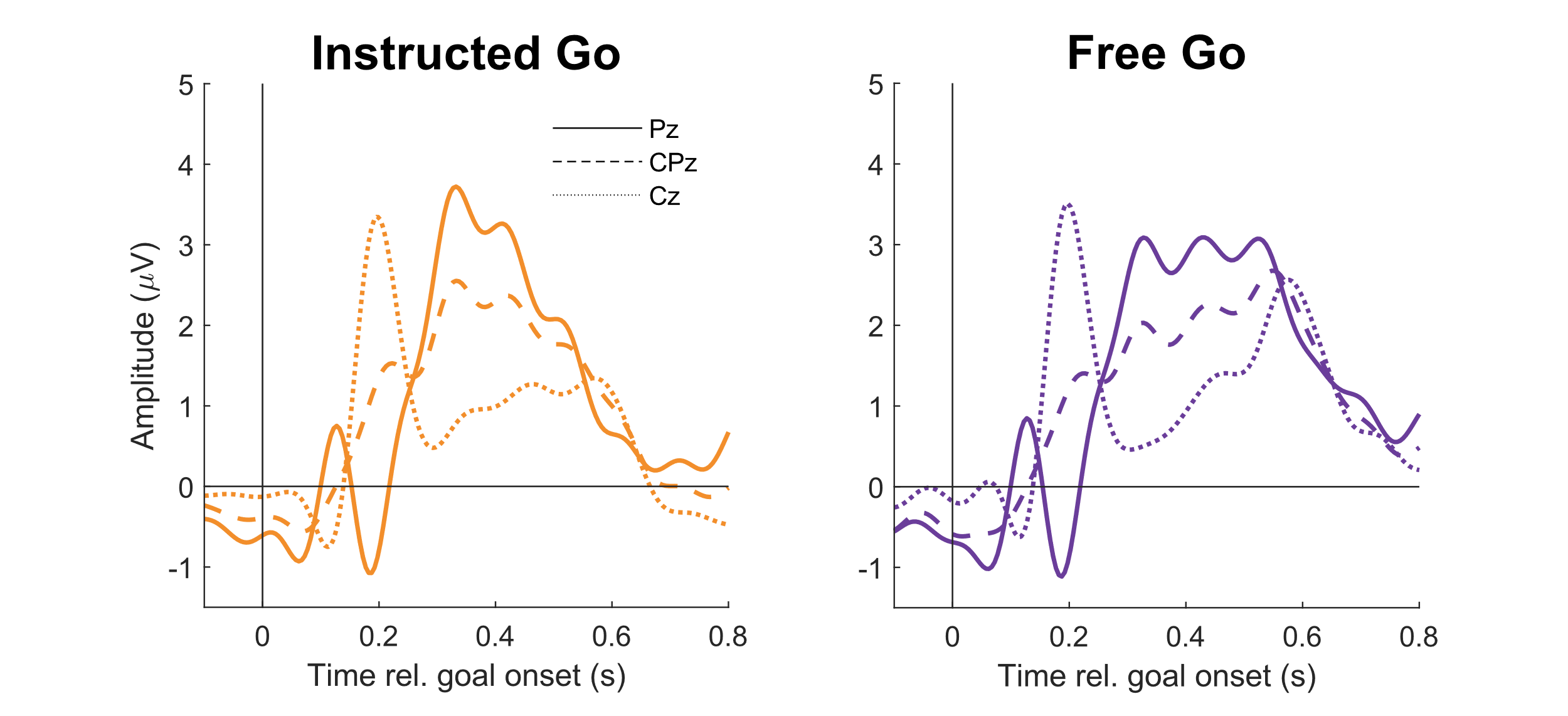

**Supplementary Figure 5.** P300 component following goal onset at centroparietal electrodes. The P300 was similar for Free and Instructed (Inst) Go conditions. EEG signals were lowpass filtered at 11 Hz and baseline corrected ([-0.2,0] relative to goal onset), averaged within and then across participants.

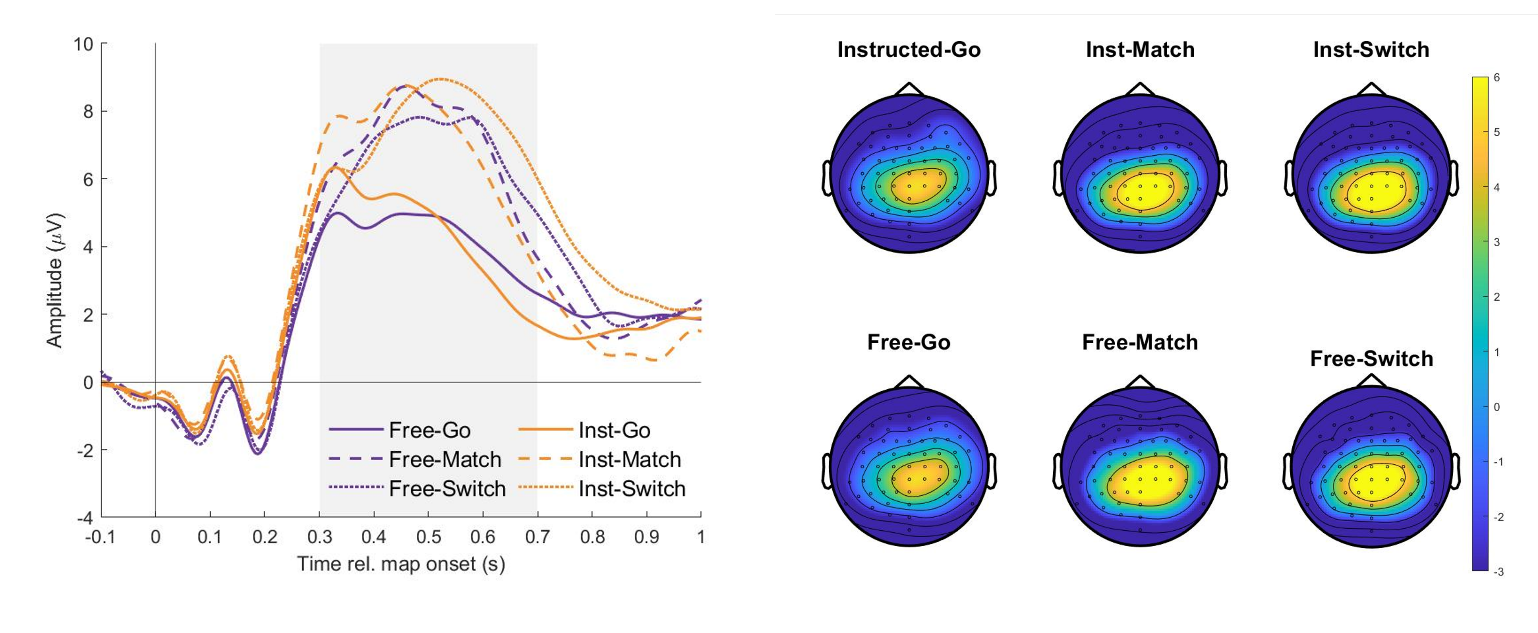

**Supplementary Figure 6.** P300 component following map onset is greater for all override conditions (Match, Switch) compared to Go conditions for both Free and Instructed contexts. Left: event-related potential at electrode Pz for all conditions (preprocessing same as in Fig, S4). Right: topoplots showing distribution of P300 component in the period [0.3, 0.7] s relative to map onset (shaded region in left panel).

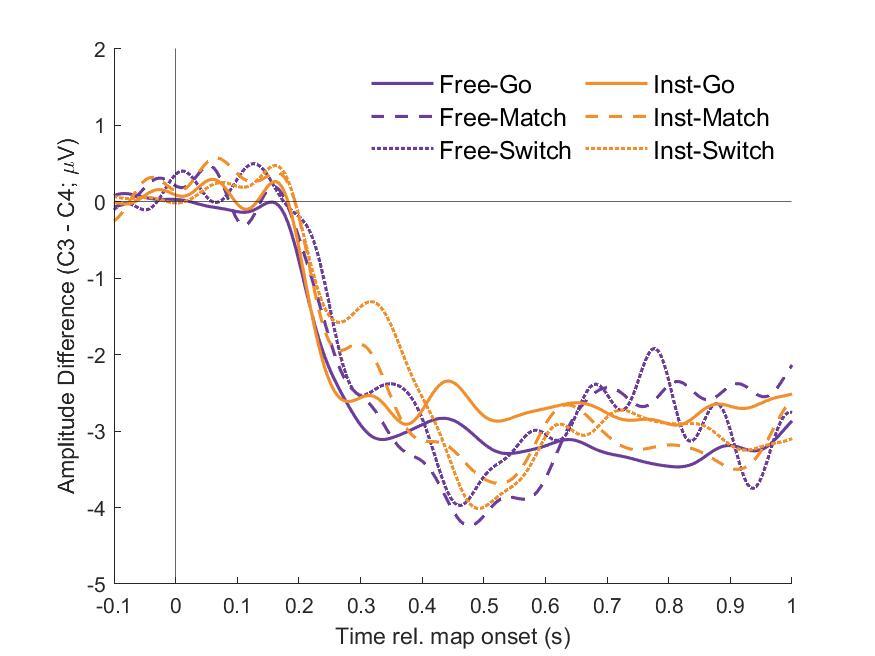

**Supplementary Figure 7.** Lateralized Readiness Potential (LRP) aligned to map onset (as in Furstenberg et al., 2023). EEG was preprocessed as in Fig. S4, and the LRP was defined as the difference in amplitude between electrodes C3 and C4 (over left and right motor cortex respectively, note that participants made all button-presses with their right hand).

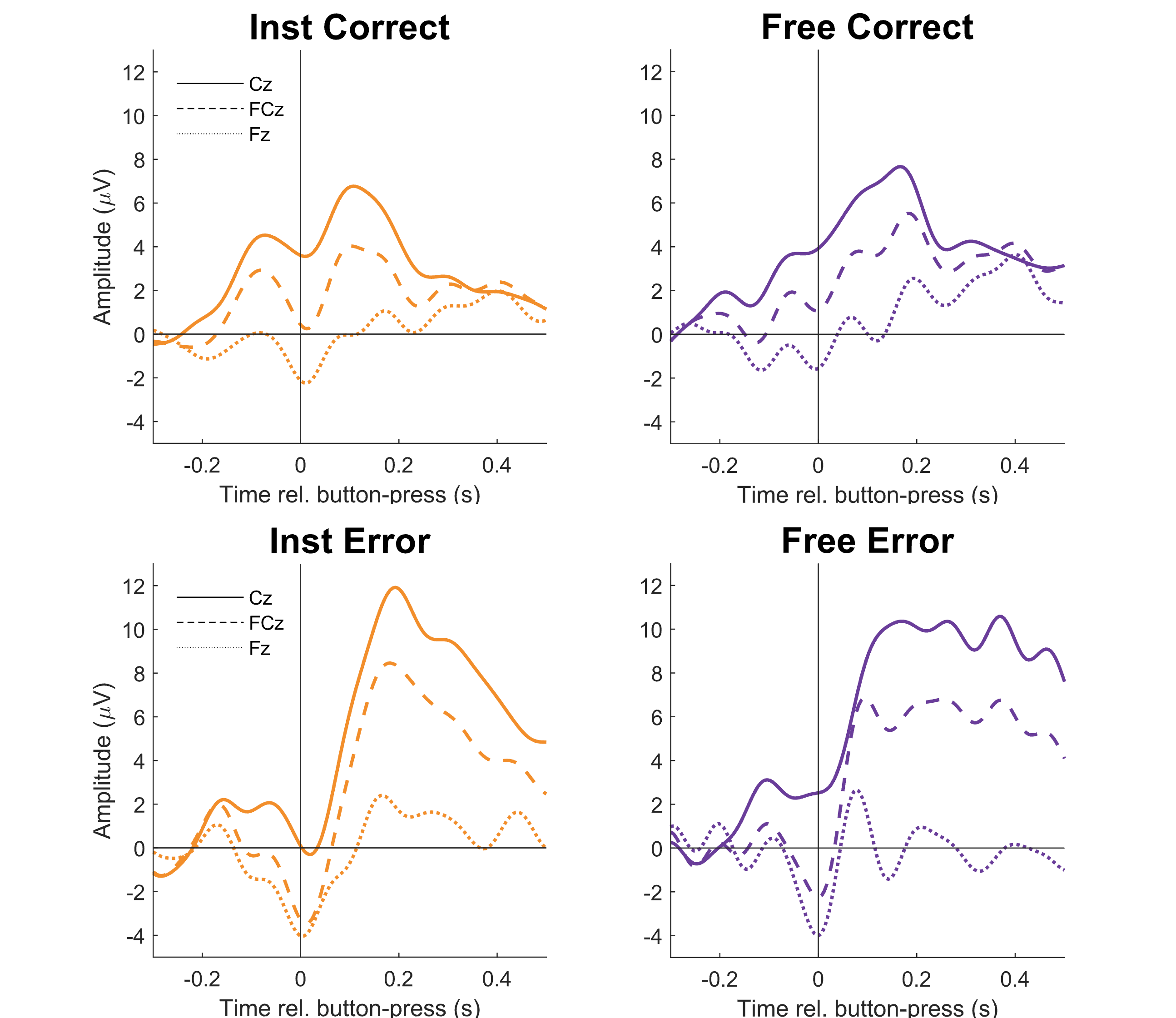

**Supplementary Figure 8.** Error-related negativity (ERN) and Correct-related negativity (CRN) aligned to button-press at frontocentral electrodes. EEG was preprocessed as in Fig S4, except we used a baseline of [-0.2, -0.1] relative to button-press to account for earlier emergence of the ERN. Negativities around button-press were similar for Free and Instructed conditions at Fz but showed differences for more central electrodes.

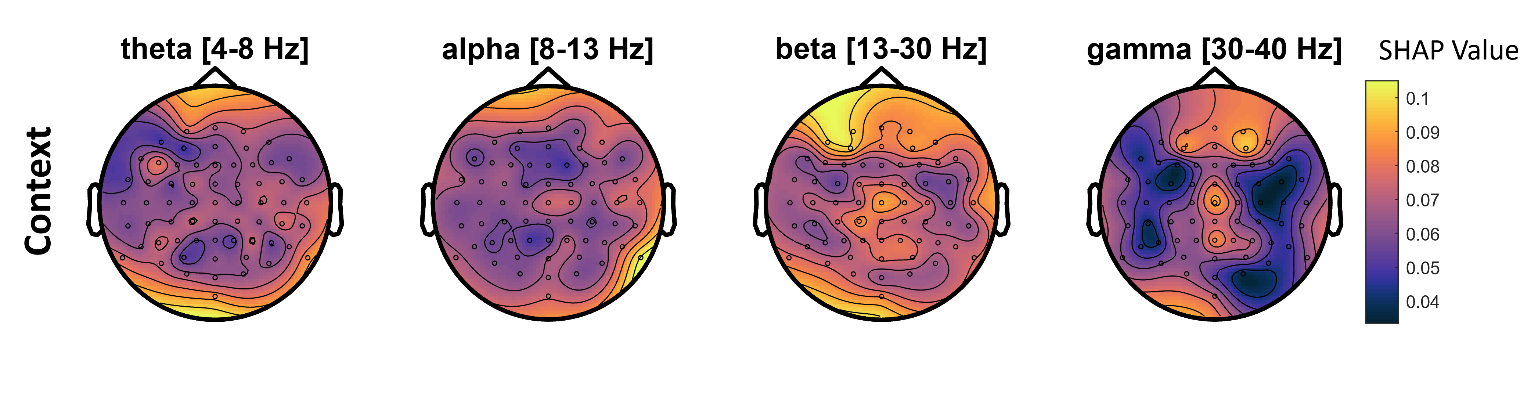

**Supplementary Figure 9.** Absolute SHAP value-based importance maps by frequency band for context decoders during the period [0.2,0.7] s relative to goal onset (see Fig 4C). For this analysis, we extracted unsigned SHAP values for each decoding feature (frequency-specific) and averaged within frequency bands to create these importance maps.

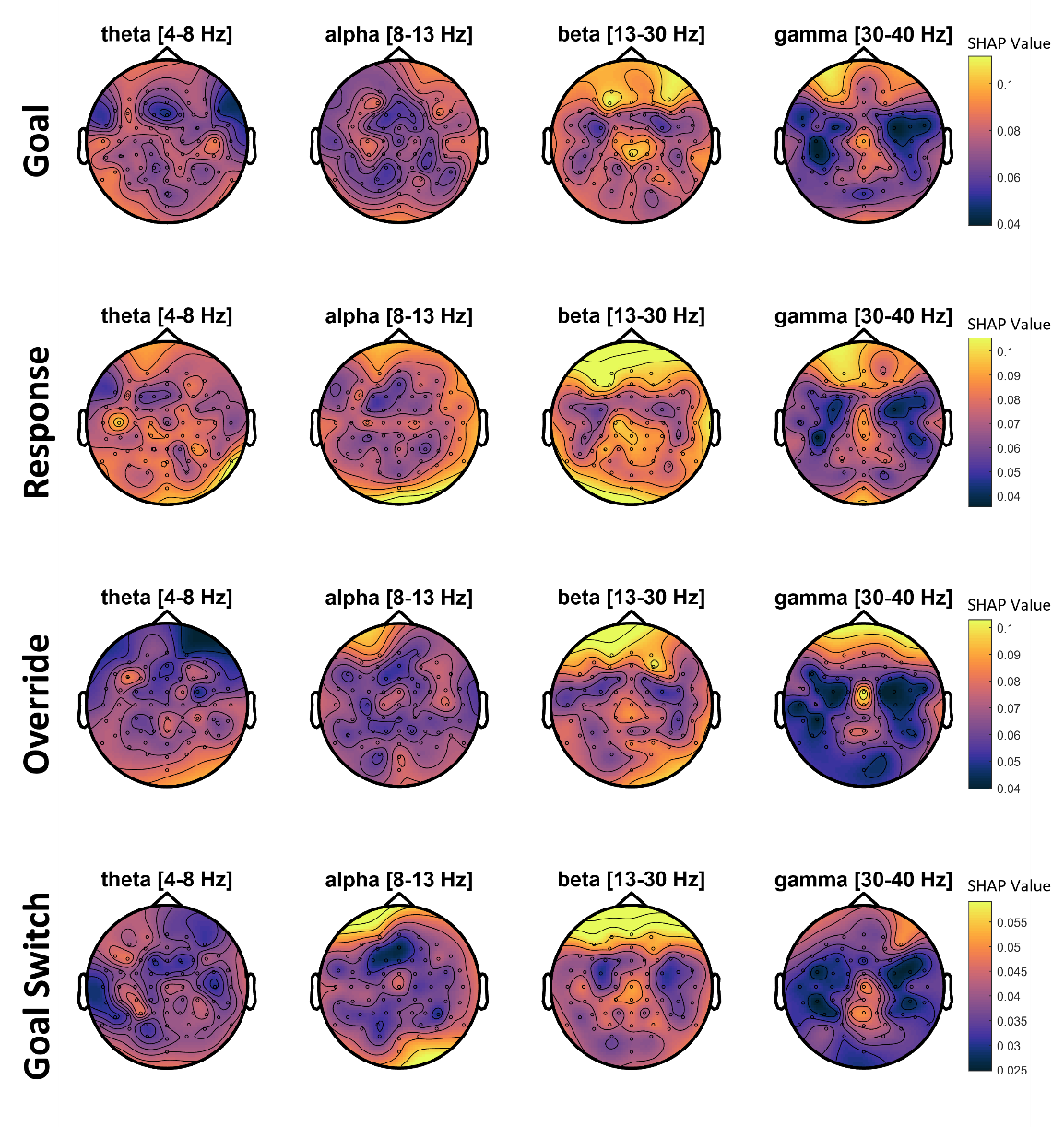

**Supplementary Figure 10.** Absolute SHAP value-based importance maps by frequency bands for Goal, Response, Override, and Goal Switch decoders in the period [-0.5, 0] s relative to button-press (see Fig 4D). Importance maps were defined as in Fig S8.
